# Multimodal phenotyping defines variant-to-function maps for RBM20 in dilated cardiomyopathy

**DOI:** 10.64898/2026.06.10.730167

**Authors:** Kai Fenzl, Linda H. Müller, Julia Kornienko, Maria Majchrowska, Nils Eikmeier, Benjamin Wehnert, Edukondalu Mullapudi, Brendan J. Floyd, Brian Clarke, Kristian Schweimer, Janosch Hennig, Daniel Schraivogel, Victoria N. Parikh, Matthias Wilmanns, Lars M. Steinmetz

**Affiliations:** Genome Biology Unit, European Molecular Biology Laboratory (EMBL); Heidelberg, Germany; DZHK (German Centre for Cardiovascular Research); partner site Heidelberg/Mannheim, Germany; Faculty of Biosciences; Heidelberg University, Heidelberg, Germany; Faculty of Chemistry and Earth Sciences; Heidelberg University, Heidelberg, Germany; Faculty of Engineering Sciences; Heidelberg University, Heidelberg, Germany; Structure Biology Unit, European Molecular Biology Laboratory (EMBL); Hamburg, Germany; Stanford Center for Inherited Cardiovascular Disease and Department of Medicine, Stanford School of Medicine; CA, USA; Division of Computational Genomics and Systems Genetics, German Cancer Research Center (DKFZ); Heidelberg, Germany; Department of Biochemistry IV - Biophysical Chemistry, University of Bayreuth; Bayreuth, Germany; Molecular Systems Biology Unit, European Molecular Biology Laboratory (EMBL); Heidelberg, Germany; Department of Genetics, Stanford University School of Medicine; CA, USA; Max Delbrück Center for Molecular Medicine; Berlin, Germany; Institute for Clinical Chemistry and Laboratory Medicine, Technische Universität Dresden; Dresden, Germany; Department of Proteomics and Signal Transduction, Max Planck Institute of Biochemistry; Martinsried, Germany; BioNTech SE; Mainz, Germany

## Abstract

Multiplex assays of variant effects have linked thousands of genotypes to fitness effects, yet we lack profound understanding of how variants impact molecular phenotypes. Here, we introduce a deep mutational scanning framework that quantifies disease-determining molecular phenotypes in human cells, allowing readouts of protein localization and splicing regulatory function at scale. Applied to the dilated cardiomyopathy (DCM)-associated protein RBM20, we profiled ∼4,300 amino acid substitutions across disease-linked protein domains. Complemented by structure-function investigations of RBM20 bound to its nuclear import receptor TNPO3, we discover new variant hotspots affecting protein function. Finally, we systematically probed nuclear relocalization to identify variants that may be amenable to this therapeutic strategy. Together, we create comprehensive variant-to-function maps that predict variant impact, enhance clinical interpretation, and stratify RBM20-mediated DCM into mechanistically distinct therapeutic classes.

## Main Text

Precision medicine requires mechanistic links from genetic variants to disease phenotypes. Yet for most missense variants, causality and mechanism remain unresolved, and variants of uncertain significance (VUS) continue to outpace pathogenic or benign annotations^1^. Deep mutational scanning (DMS), paired with multiplexed assays of variant effects (MAVEs), enables assaying single amino acid substitutions at scale. Integrating multiple complementary readouts was recently shown to improve variant interpretation^2–4^. Here, we applied such a combined pooled approach to systematically assay ∼4,300 single amino acid substitutions across three disease-associated domains of RBM20 (RNA-binding motif protein 20), an alternative splicing regulator implicated in familial dilated cardiomyopathy (DCM).

DCM is a leading cause of heart failure and heart transplantation^5^. All ClinVar-annotated disease-causing missense variants in *RBM20* map to the protein’s RNA Recognition Motif (RRM), arginine and serine (RS)-rich, or glutamate (E)-rich domains^6^. The PRSRSP stretch, a hotspot for DCM-associated mutations located in the intrinsically disordered RS-rich domain, is implicated in RBM20 nuclear import via direct interaction with Transportin-3 (TNPO3)^7^. The folded RRM domain was shown to be required for RNA binding and splicing function, and the intrinsically disordered E-rich domain to mediate protein-protein interactions and protein stability^8^. Thus, depending on the variant within RBM20, its pathogenicity is based on distinct mechanisms, including missplicing of RBM20 targets, such as *TTN* transcripts, with or without mislocalization. It has been demonstrated that RBM20 mislocalization caused by PRSRSP-stretch missense mutations can drive a toxic, neomorphic gain-of-function phenotype beyond splicing loss-of-function^9,10^. These variants are associated with a more severe disease prognosis than variants that drive a splicing dysfunction alone^6,11^. TNPO3 overexpression was shown to restore nuclear localization, splicing, and cardiac function in mice with two pathogenic variants located in the PRSRSP-stretch^12^, raising the prospect of disease rescue via RBM20 relocalization. However, it is not well understood whether missense variants outside the PRSRSP-stretch can also lead to mislocalization, and if so, whether therapeutic nuclear relocalization can be generalized to those. Our study provides comprehensive variant-to-function maps aiming to improve clinical variant interpretation and guide mechanism-tailored therapeutic strategies, including pooled testing of re-localization as a functional rescue approach.

To systematically map variant effects on localization and splicing regulation, saturation mutagenesis libraries were designed for the disease-associated RS-rich, RRM, and E-rich domains of RBM20 (**Fig. 1A**). We exchanged each wild-type (WT) residue with each of the other 19 amino acids. The mutagenesis libraries were cloned into lentiviral plasmids, and HeLa Kyoto cells were transduced to generate stable, single-variant cell pools, which naturally lack endogenous expression of the cardiac RBM20 splicing factor^13^. A balanced variant representation was achieved (**Fig. S1A**). Image-enabled cell sorting (ICS), which combines fluorescence imaging at subcellular resolution with high-throughput cell sorting^14^, was applied to isolate cells with nuclear (WT-like) or cytoplasmic (mislocalized) EGFP-RBM20 variants. The separation was achieved based on EGFP co-localization with the nuclear dye DRAQ5^7^ (**Fig. 1B**, **Fig. S1B-D**). Amplicon sequencing on genomic DNA from sorted cells was performed to quantify variant representation. To read out the splicing function in a pooled format, we adapted a *Ttn* splicing reporter mini gene^15^. The *Ttn* mini gene reports on the relative abundance of N2B- and N2A-type *Ttn* isoforms by the ratio of two fluorophores, enabling a splicing-dependent cell sort (**Fig. 1B, Fig. S1E, F**). Benchmarking confirmed that this assay captures the known difference between RBM20-WT and the mislocalizing, splicing-deficient P633L and R634Q variants^7^, as well as a Ser635FrameShift loss-of-function variant^16^ (**Fig. S1G**).

**Fig. 1:**
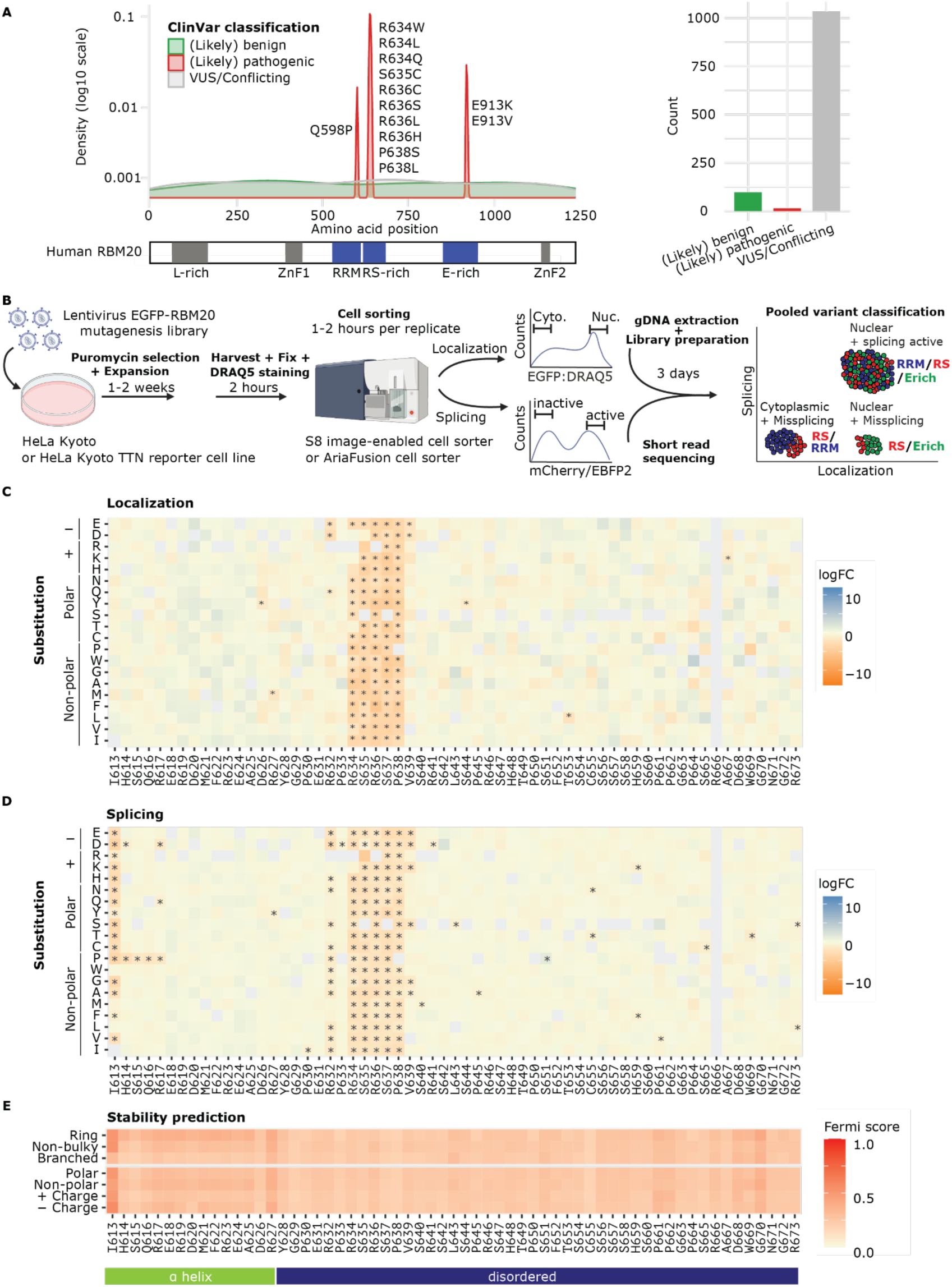
Multimodal RS-rich domain screening reveals variants affecting RBM20 function. **(A)** Left: Density plot of RBM20 missense variant annotations from ClinVar (August 2025). Only (likely) pathogenic missense variants are labeled. RBM20 domains with (likely) pathogenic variants are highlighted in blue. All reported variants arise by a single nucleotide exchange. Right: Number of deposited *RBM20* missense variants for each classification. **(B)** Experimental set-up for pooled localization and splicing screening. **(C-D)** Variant effect maps for the RS-rich domain. Gray tiles indicate missing variants or WT residues. Heatmaps show results from at least two replicates, stars indicate significant single variant hits (adj. p < 0.05). Effect sizes were normalized to 10 benign-predicted variants (see materials and methods). **(E)** Protein fold stability prediction using RaSP^17^, mean effect sizes are grouped by amino acids of similar chemical properties. Predicted secondary structures (AlphaFold^25^) are marked.

### Variant effect maps of the RS-rich domain

We first analyzed the RS-rich domain, which includes several reported mislocalizing variants within the RSRSP stretch (residues 634-638)^6–9^. All 95 possible RSRSP hotspot variants induced mislocalization, with the exception of six variants not reaching significance (**Fig. 1C**). All known pathogenic RS-rich domain variants (RS variants) were detected as significantly mislocalized in our screen, except for the mildly mislocalizing P633L variant. Additionally, flanking positions R632 and V639 impacted nuclear localization when substituted with negatively charged residues, extending the already known RSRSP hotspot. These data allow to rank all single amino acid substitutions by mislocalization severity. Notably, the top mislocalizing hits require at least two nucleotide exchanges in one codon and are therefore not included in known pathogenic RS variants, which arise exclusively from single nucleotide exchanges (**Table S1, Fig. S1H**).

As anticipated, all mislocalizing variants within the RSRSP hotspot were splicing-deficient, including several substitutions at the flanking positions R632 and V639 (**Fig. 1D**). In addition, substitutions at I613 decreased splicing activity without affecting nuclear localization, and introducing proline at adjacent residues 614-617 likewise caused missplicing (**Fig. 1D**). These residues map to an α-helix, which forms upon RNA binding, and is required for RNA recognition^16^. We applied Rapid Stability Prediction (RaSP)^17^ to indicate how each substitution may affect protein fold stability. Disruptions at position 613 were predicted to impair stability, whereas all other RS variants did not indicate a substantial effect (**Fig. 1E**), in line with our splicing data. Together, these results provide a comprehensive variant effect map of the RS-rich domain that validated the known RSRSP hotspot and identified previously uncharacterized variants affecting RBM20’s localization and/or splicing function (**Table S1**).

### Variant effect maps of the RRM domain

We next investigated the RRM domain of RBM20 - a folded, functional domain abundant across the human proteome, particularly in RNA-binding proteins^18^. The pooled localization and splicing screens revealed a multitude of residues highly sensitive to substitution, with nearly all changes at these sites causing significant mislocalization and/or missplicing (**Fig. 2A, B**). As expected, detected mislocalization hotspots were also prominent hotspots for splicing-deficiency. Variants within regions with secondary structures were more prone to cause damage than variants in unstructured regions (**Fig. S2A, B**). In addition, functional localization and splicing scores correlated with protein fold stability predictions according to RaSP (Pearson r = 0.52, p < 0.0001 for localization; Pearson r = 0.43, p < 0.0001 for splicing; **Fig. 2C**). Damaging variants, including loss- and gain-of-function variants, typically included substitutions to proline, or substitutions of branched (isoleucine, leucine, valine) towards non-branched residues within secondary-structure elements (**Fig. 2A, B**), pointing towards misfolding of the RRM domain as cause of the functionality loss.

**Fig. 2:**
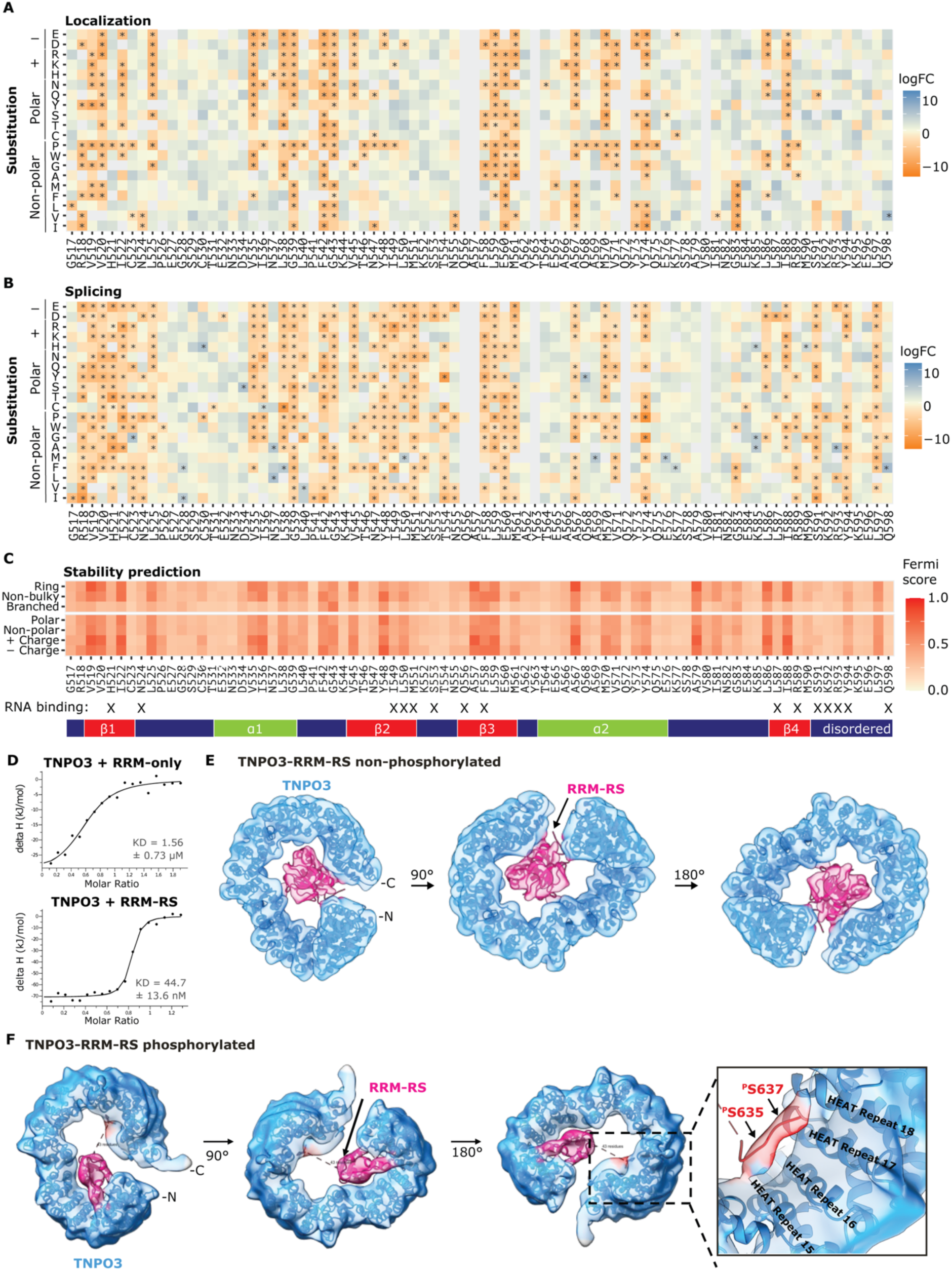
Multimodal RRM domain screening and the interaction with TNPO3. **(A-B)** Variant effect maps for the RRM domain. Gray tiles indicate missing variants or WT residues. Heatmaps show results from at least two replicates, stars indicate significant single variant hits (adj. p < 0.05). Effect sizes were normalized to 10 benign-predicted variants (see materials and methods). **(C)** Protein fold stability prediction using RaSP^17^, mean effect sizes are grouped by amino acids of similar chemical properties. Predicted secondary structures (AlphaFold^25^), and interaction sites with RNA^19^ are marked. **(D)** Isothermal titration calorimetry (ITC) measurements of TNPO3 with RRM alone (top) or RRM-RS constructs (bottom). **(E-F)** Cryo-EM reconstructions of TNPO3 in complex with the unphosphorylated (E) or phosphorylated (F) RRM-RS domain of RBM20.

In addition to mislocalizing and missplicing variants, we identified several splicing-deficient variants that were not detected as mislocalizing (**Fig. 2B**). These were enriched at positions implicated in RNA binding^19^ (H521, N524, I549-M551, S591, Y594), including the second β-sheet, suggesting loss of RNA binding as the leading cause of the observed effect. In line with a previous study showing compromised RNA-binding upon substitutions in the RRM domain of murine RBM20 (H523A, F560A, and Y596A^19^), our screen identified the respective human H521A, F558A, and Y594A variants as significantly splicing-impaired (**Fig. S2C**). Further, the missplicing I536T variant (orthologous to I538T in mice) was previously shown to disrupt splicing of *Ttn* and other endogenously expressed targets, including *Ldb3, Camk2d*, and *Ryr2,* in mice^20^.

Together, our multimodal phenotyping of the RRM domain identified previously unknown residues crucial for RBM20 nuclear import and distinguished mislocalizing from nuclear splicing-deficient variants. Previous studies showed that the RRM-domain variant V535I^21^ and the likely pathogenic ClinVar-reported variant Q598P^22^ did not affect the protein’s localization, which is in line with our data (**Fig. 2A**). A full RRM deletion has been reported to either partially disrupt^13^ or not affect localization^21^. RRM missense variants could disrupt the interaction with the nuclear import receptor TNPO3, which was previously only linked to the structurally nearby RS-rich domain.

### Structure of RRM-RS domain with TNPO3

To test the participation of the RRM domain in TNPO3 binding, we first performed isothermal titration calorimetry (ITC) with a purified isolated RRM domain or combined RRM-RS construct (both, RRM- and RS-rich domains). The isolated RRM domain bound TNPO3 with a micromolar dissociation constant (KD), while the presence of the RS-rich domain enhanced the affinity by 100-fold (**Fig. 2D, Fig. S2D**), indicating cooperative binding.

RBM20’s RSRSP stretch is known to be constitutively phosphorylated and crucial for efficient nuclear transport^15^. However, our RS-rich domain localization screen indicated that phosphomimetic variants (aspartate, glutamate) at S635 and S637 cause mislocalization (**Fig. 1C**), in agreement with already published data^23^. To understand the structural basis of the interaction between TNPO3 and RBM20 as well as the impact of phosphorylation, we determined Cryo-EM structures of the RRM-RS/TNPO3 complex with both phosphorylated and unphosphorylated RRM-RS constructs (**Fig. 2E, F**). Serine phosphorylation, as well as TNPO3 binding before and after RS-rich domain phosphorylation, were confirmed by NMR (**Fig. S2E-G**). Both Cryo-EM density maps revealed TNPO3 as an α-helical arch-like structure, which consists of 20 pairs of HEAT repeats^24^. TNPO3 was bound to a flexible and less defined RRM-RS construct, preventing a direct interpretation of the binding interface from the experimental maps alone (resolution of >6 Å). Therefore, we used AlphaFold 3 (AF3)^25^ to model and validate the precise binding mode for each state (see Materials and Methods). The high-confidence AlphaFold model suggests fundamentally different binding conformations: (i) In the unphosphorylated state, the RRM domain is bound to the C-terminus of TNPO3, whereas no binding for the RS domain was detectable in the density map; (ii) In the phosphorylated state, the RS-rich domain engaged the C-terminal region of TNPO3 and the RRM domain bound to the opposite N-terminus (**Fig. 2E, F**). The binding in the phosphorylated state resembles the solved structure of TNPO3 in complex with the RRM2-RS domain of the alternative splicing factor ASF/SF2^24^ and indicates close vicinity of the R-pS-R-pS-R motif to TNPO3’s HEAT repeats 14-17 (**Table S2**).

In summary, our data show that while phosphorylation of RBM20 acts as a molecular switch that dictates RBM20’s binding orientation to TNPO3, the *in vitro* interaction of RBM20 and TNPO3 is phosphorylation-independent and involves the RRM domain.

### Variant effect maps of the E-rich domain

Lastly, we screened the intrinsically disordered E-rich domain. We did not identify hotspots or single residues that significantly altered nuclear RBM20 localization (**Fig. 3A**), as expected from published single variant studies^21^. However, the splicing screen revealed a consecutive stretch of splicing-deficient variants (residues 905-916, **Fig. 3B**). Notably, the only two likely pathogenic ClinVar-annotated variants within the E-rich domain, E913K (previously linked to abnormal isoform ratios of *TTN*^8^) and E913V, map to this hotspot. Substituting residues V909, V911, and V914 with another branched amino acid did not affect the splicing activity, and negatively charged residues E906 and E907 tolerated substitution with negatively charged aspartate. Since missense variants within the hotspot did not mislocalize, we hypothesize that these residues contribute to protein stability, as previously described^8^, or protein-protein interactions essential for spliceosome assembly or function. However, only few variants of the disordered E-rich domain indicated misfolding in the RaSP protein fold stability prediction (**Fig. 3C**). Thus, either this region may act as a protein-protein interaction hub required for splicing, or the destabilization is misfolding-independent.

**Fig. 3:**
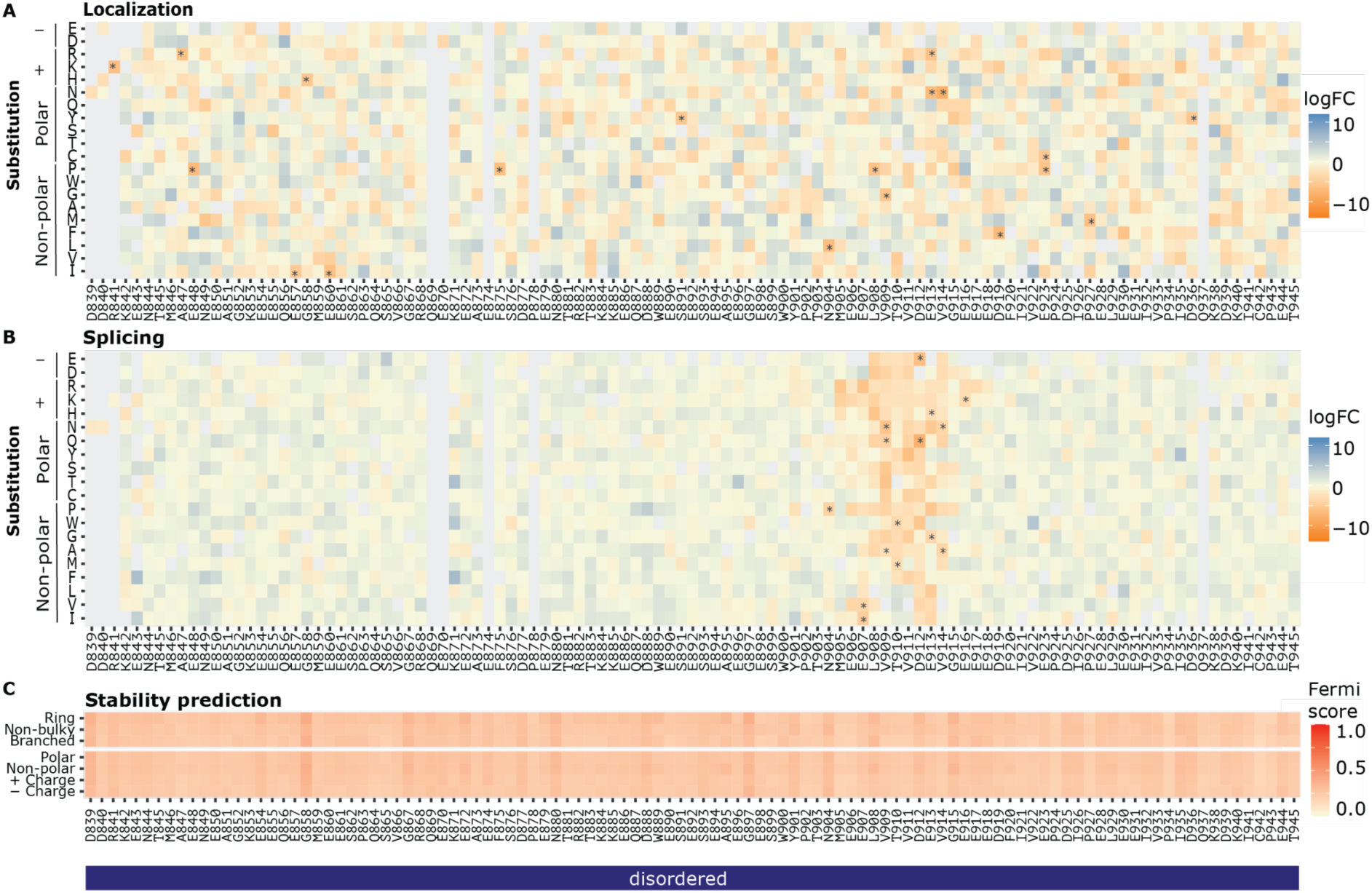
Deep mutational scanning of the E-rich domain. **(A-B)** Deep mutational scanning results for the E-rich domain mutagenesis library depicted as heat maps. Gray tiles indicate variants with missing data or WT residues. Heatmaps show results from at least two replicates, stars indicate significant single variant hits (adj. p < 0.05). Effect sizes were normalized to 10 benign-predicted variants (see materials and methods). **(C)** Protein fold stability prediction using RaSP^17^, mean effect sizes are grouped by amino acids of similar chemical properties. Secondary structure prediction (AlphaFold^25^) is marked.

### Functional validation of single variants

We tested six variants spanning all screened domains to individually confirm localization and splicing effects. The RRM V571F variant, which was expected to mislocalize based on our localization screening data, showed severe mislocalization and cytoplasmic granule formation (**Fig. S3A, B**). Three out of the five tested variants that did not meet the significance threshold in the localization screen displayed mild mislocalization in the arrayed validation (RS: I613P, S635D; RRM: M551I), indicating a screen sensitivity limit for subtle or partial mislocalization. Nonetheless, all three caused loss-of-splicing, both in the arrayed validation and screen, classifying them as damaging (**Fig. S3C**). This highlights the advantage of multimodal phenotyping for advanced variant assessment. Overall, agreement between the pooled screen and arrayed validation was strong for both localization (Pearson r = 0.90) and splicing (r = 0.92) (**Fig. S3D**), and effects identified as significant in the pooled screen were confirmed during validation, underlining the confidence in functionally defective hits.

Furthermore, our RBM20 localization and splicing scores are in agreement with various published single variant-focused studies across more physiologically relevant model systems, including patients, mouse models, and iPSC-derived cardiomyocytes (detailed overview provided in **Table S3**).

### Systematic evaluation of nuclear re-localization

Relocalization of RBM20 is a potential precision medicine approach for patients with mislocalizing RBM20 variants^12^. However, it is unknown whether all mislocalized RRM and RS-rich domain RBM20 variants retain their splicing function after relocalization, as previously shown for single RSRSP-stretch variants^7^. We demonstrated that N-terminal tagging of RBM20-R634Q with a nuclear localization signal (NLS) in human iPSC-derived cardiomyocytes led to a similar degree of splicing rescue as we observed upon TNPO3 overexpression *in vitro* and *in vivo*^7,12^. Therefore, to probe splicing rescue at scale, we applied a similar approach, fusing all RS and RRM variants to an N-terminal NLS. ICS measurements and fluorescence microscopy verified nuclear localization of variant libraries (**Fig. 4A**). Within the RS-rich domain, the screen indicated robust splicing rescue for RSRSP hotspot variants (residues 634-638) as well as for the newly identified mislocalizing variants at flanking positions R632/V639 (**Fig. 4B, Fig. S4A, Table S4**). Consistent with defects in RNA binding and/or fold integrity, the majority of RRM variants showed less pronounced rescue after NLS-mediated relocalization, with some exceptions (**Fig. 4B, Fig. S4B**).

**Fig. 4:**
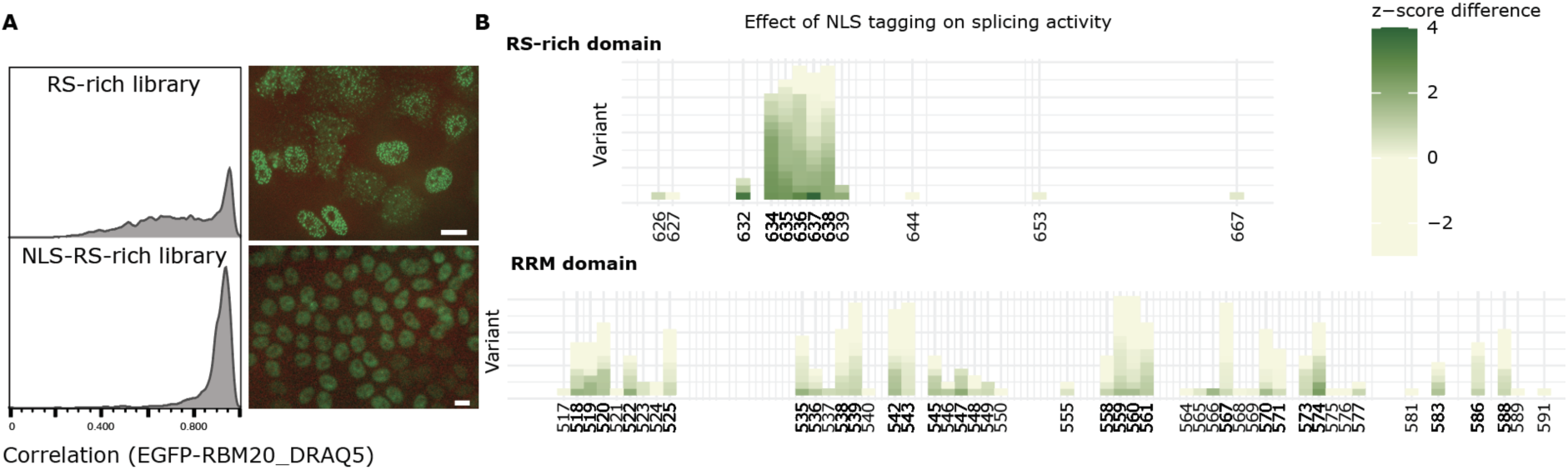
Systematic evaluation of nuclear re-localization. **(A)** ICS EGFP:DRAQ5 correlation plot for RS-rich domain HeLa Kyoto cell pools with or without NLS-tag, and representative microscopy images. Scale bars indicate 10 µm. **(B)** Effects of NLS-tagging on splicing activity for mislocalizing (localization screen adj. p < 0.05) RS-rich (top) and RRM domain (bottom) variants. Depicted as the difference in z-scores of logFCs between NLS- and not-tagged splicing screens. Green indicates rescue via nuclear re-localization. Heatmaps show results from at least two replicates.

### Linking functional splicing scores to clinical pathogenicity

To evaluate the consistency of our RBM20 variant effect catalog with clinical data, we integrated splicing scores (reflecting mislocalization and loss of splicing, **Fig. 1D**, **Fig. 2B**, **Fig. 3B**) with curated clinical annotations (**Table S5**, see Materials and Methods). We first assessed how well our functional scores reflect the pathogenicity of known disease-associated variants, and observed a clear separation between likely pathogenic and likely benign variants (p < 0.0001; **Fig. 5A**). The pathogenic variant in our screen that had a functional score most similar to WT-like variants was P633L, which was previously shown to only mildly mislocalize and feature residual splicing activity^7^ (**Fig. S1G**). This highlights the accuracy of pathogenicity classification and ranking based on our functional scores. Interestingly, several reported VUS (e.g., V535F) demonstrated scores similar to those of known pathogenic variants (**Fig. 5A**), suggesting their potential pathogenicity.

**Fig. 5:**
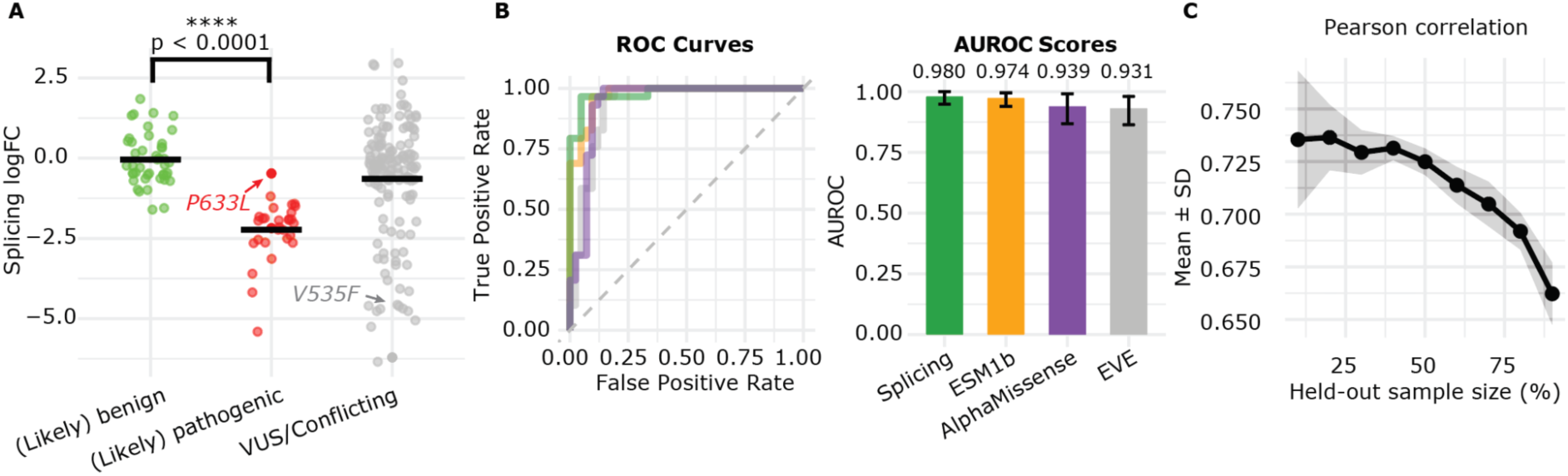
Clinical relevance of variant classification. **(A)** Measured splicing effect sizes for annotated (likely) benign, (likely) pathogenic and VUS/Conflicting variants based on ClinVar annotations and internal clinical curation (see materials and methods). Statistical significance assessed by two-sided t-test. **(B)** ROC curves (left) and corresponding AUROC values (right) for score comparisons in regards to classification of annotated (likely) benign and (likely) pathogenic RBM20 variants. Only variants with complete data across all scores were included in the analysis: 42 (likely) benign and 29 (likely) pathogenic variants (see **Table S5**). Bootstrapping with 1000 iterations was performed and 95% confidence intervals are indicated. **(C)** Random forest models were trained to predict splicing logFCs using randomly sampled variant subsets from 10% to 90%. For each level, the variant training set was randomly sampled 10 times with different seeds. Reported values are the mean and standard deviation across repetitions. Pearson correlation of observed and predicted splicing logFCs.

We next benchmarked variant effect predictors (VEPs), including AlphaMissense^26^, EVE^27^, and ESM1b^28^, against our functional scores (**Fig. S5A**). On the curated RBM20 annotation set (42 likely benign vs. 29 likely pathogenic variants), our functional scores achieved a higher AUROC (0.980) and exceeded all tested VEPs (0.931–0.974, **Fig. 5B**). This is in line with prior observations that DMS surpasses VEPs in correctly categorizing annotated benign and pathogenic variants^29^. We observed that VEP scores predict a higher number of variants as pathogenic compared to our experimental scores (**Fig. S5B**), thus potentially overestimating this number. Importantly, and in contrast to *in silico* predictions, our study provides mechanistic insights (**Table S1**). E.g., the V535F variant (VUS) mislocalizes (**Fig. 2A**) and thus can be classified as gain-of-function with severe disease prognosis. However, V535F did not show splicing rescue after nuclear relocalization (**Fig. S4B, Table S4**), highlighting that any relocalization approach would not rescue isoform imbalances. In total, we detected 321 mislocalizing gain-of-function variants (logFC < -0.5 & adj. p < 0.05 for splicing and localization) and 147 loss-of-function variants affecting splicing without mislocalization (logFC < -0.5 & adj. p < 0.05 for splicing & logFC ≥ 0 for localization) (**Fig. S5C, Table S1**). These data highlight the advantage of integrating multiple complementary readouts to achieve more accurate variant interpretation and thus suggest best-suited therapeutic approaches for each variant.

Lastly, to explore scalability and potential cost preservation via coupling DMS to machine learning, we performed downsizing simulations. We randomly masked 10-90% of variants, trained random-forest models using localization scores and 46 publicly available features (e.g., amino acid properties, VEP scores, conservation; **Table S6**), and predicted held-out splicing scores. Holding out up to 70% of variants retained strong performance (mean Pearson r > 0.7; AUROC > 0.95 for clinically-annotated variants; **Fig. 5C**, **Fig. S5D, E**), demonstrating the possibility to shrink DMS size and thus the associated experimental cost and time.

## Discussion

By leveraging image-enabled cell sorting (ICS) for assessing localization and splicing activity, we mapped effects for over 4,300 RBM20 amino acid substitutions (92% coverage of RS-rich, RRM, and E-rich domains), including 154 reported VUS. We found that 9% of screened variants significantly altered protein localization, whereas 15% led to missplicing. In the RS-rich domain, the vast majority of substitutions in the known mislocalization hotspot (residues 634-638) showed impaired nuclear localization, which our screen expanded to substitutions of flanking positions R632/V639 toward negatively charged residues. We identified a novel potentially clinically-relevant missplicing hotspot centered around position 613, part of an α-helical element required for RNA recognition^16^ but not nuclear localization.

Similar to the RS-rich domain, RRM domain variants suggest two pathomechanisms based on our assay. First, substitutions disrupting secondary structures led to both mislocalization and missplicing. Second, clusters of nuclear-localized, splicing-defective variants were located at RNA-binding residues, indicating that loss of RNA recognition is the leading cause of the splicing defect. In addition, we showed that the isolated RRM domain alone binds the nuclear import receptor TNPO3 and that adding the RS-rich domain increases the binding affinity by 100-fold, indicating cooperative recognition and interaction to achieve nuclear import. Several RRM domain variants may affect nuclear localization due to their proximity to the RS-rich domain. Therefore, structural changes in the RRM could disrupt its interaction with the nuclear import receptor TNPO3, previously linked only to the RS-rich domain.

Our single variant validations indicate that also RRM substitutions show cytoplasmic granule formation. This was surprising because previous full RRM domain knockout studies indicated no mislocalization as well as no DCM phenotype in rodents^21^. Our mutagenesis screen thereby shows that such knockout models can mask a gain of function, as also seen for RS-rich domain or full RBM20 knockouts versus single amino acid substitution RS models^7,9,13,21^. Interestingly, there have been no mislocalizing RRM variants annotated as pathogenic to date^11^, raising the question whether these variants may be lethal. However, eleven mislocalizing RRM variants are listed in gnomAD^30^, each with a rare allele frequency of ≤ 7.7×10⁻⁶. These include the VUS

M561T variant we identified as gain-of-function, which has been reported by three submissions linked to cardiovascular phenotypes or specifically DCM^31^. Substitutions in the C-terminal, intrinsically disordered E-rich domain showed largely unaffected localization, however, with a splicing-deficient region (905-916) including the reported DCM-causing E913K variant.

We observed that both phosphorylated and non-phosphorylated RRM-RS constructs bound TNPO3 *in vitro*, albeit at distinct sites, hinting towards a phosphorylation-independent TNPO3-mediated nuclear import of RBM20. TNPO3 binding of another RS domain from the alternative splicing factor ASF/SF2 was phosphorylation-dependent^32^, in agreement with the resolved RSRSP stretch in the phosphorylated state. Furthermore, the positioning of the RRM domain within TNPO3 matched between RBM20 and ASF/SF2^24^. While our structural and biophysical data assume a TNPO3-centric model in agreement with published data^7^, variant-specific transporter dependencies and potential roles for alternative importins could exist and were not assessed here.

To enable scalable DMS screens for further disease-associated human proteins in the future, we explored ways to reduce the costs by narrowing the assayed space from 23,000 amino acid substitutions across the full RBM20 sequence to 4,750 variants within disease-relevant domains. This prioritization, combined with technological advances and predictive model training, will lower costs and time, increase scalability, and provide deeper mechanistic insights for any protein of interest, ultimately improving targeted therapy options for patients.

Our functional scores precisely separated known benign and pathogenic RBM20 variants (AUROC 0.98), indicating the robustness of our DMS approach to classify RBM20 variants. Additionally, our approach assigns the respective pathomechanism to each defective variant, including 321 mislocalizing and 147 splicing-deficient variants without mislocalization. Such insights support stratification of RBM20-DCM patients based on the specific variant they carry, allowing for tailored, mechanism-driven therapies. One promising therapeutic avenue is nuclear relocalization, which has been shown to rescue the disease phenotype of two DCM-causing RS variants both *in vitro* and *in vivo*^7,12^. We tested the broader applicability of this approach and detected most RSRSP-hotspot and newly identified mislocalizing variants at flanking positions R632 and V639 as rescuable at the splicing level. Thus, nuclear relocalization might benefit patients with these variants.

Altogether, we propose three therapeutically relevant classes of RBM20 variants based on their underlying pathomechanisms. Class I: mislocalizing variants with splicing rescue after nuclear relocalization (predominantly RS-hotspot), which would benefit from nuclear relocalization approaches^7,12^. Class II: mislocalizing, splicing-defective variants not rescued by relocalization (primarily RRM variants), which may benefit from cytoplasmic RBM20 granule clearance^33^. Class III: nuclear, splicing-deficient variants (E-rich hotspot and RS-rich I613), for which neither relocalization nor granule clearance is suitable. For all classes, gene editing presents a viable therapeutic option^34^. These findings provide a framework for tailoring therapeutic strategies to individual pathomechanisms of single variants, advancing precision medicine for RBM20-related diseases and beyond.

## Acknowledgments

We thank Nikolay Dobrev and Jan Blaha for technical and experimental support with structural investigations; Marta Rodriguez for experimental support with image-enabled cell sorting. Special thanks go to the EMBL Genomics Core Facility for next-generation sequencing services, and Vladimir Benes and Laura Villacorta for advice and support; the EMBL Flow Cytometry Core Facility for instrument maintenance and sorting services, and Daniel Gimenes, Beata Ramasz, and Diana Ordoñez for advice and support; the EMBL Protein Expression and Purification Core Facility for instrument maintenance, and Kim Reans, Jacob Scheurich, and Karine Lapouge for advice and support; the EMBL Data Science Center for support, and Sarah Kaspar and Charles Girardot for advice on data analysis. We thank Dr. Hidehito Kuroyanagi and the lab for providing the *Ttn* minigene splicing reporter, which we adapted for our study.

## Funding

This work was funded by the EIC Pathfinder CARDIOREPAIR 101115574, Leducq Foundation Cardiac Splicing as a Therapeutic Target (CASTT; 20CVD02), Deutsche Forschungsgemeinschaft (DFG, German Research Foundation) - SFB1550 - Project ID 464424253: Collaborative Research Center 1550 (CRC1550) “Molecular Circuits of Heart Disease”, Else Kröner Fresenius Zentrum (EKFZ; grant number 16654). KF was supported by a research fellowship from the EMBL Interdisciplinary Postdoc (EIPOD) Programme under Marie Curie Cofund Actions MSCA-COFUND-FP (grant agreement 847543). LHM was supported by the Medical Science fellowship from the Bayer Foundation.

## Author contributions

KF, LHM, DS, LMS designed the study. KF, LHM, MM, NE performed the DMS experiments. KF, LHM, BW, NE analyzed the sequencing data. KF, EM performed structural analyses and predictions. JH, KS performed the NMR measurements and data analysis. LHM performed machine learning with guidance from BC and support from BW. BJF, VNP performed clinical curation of RBM20 annotations. All authors provided expert guidance and feedback on analysis and results. KF, LHM, JK, DS wrote the manuscript with input from JH, MW, and LMS, and feedback from all authors.

## Competing interests

LMS is a co-founder and shareholder of Sophia Genetics, and has received funding from AstraZeneca and Novartis. KF, JK, and LMS filed an invention disclosure describing TNPO3 and restoring nuclear localization of RBM20 variants discussed in this article (US provisional patent application 63/452 252, filed March 15, 2023). JK, KF, and LMS filed an additional invention disclosure describing an *in silico* designed binder as an alternative RBM20 transporter in young and adult mice (US provisional patent application 63/906,830, filed October 28, 2025).

## Additional Information

Supplementary Information is available (Fig. S1 - S6, Table S1-S8). Raw and processed data are available upon request. Correspondence and requests for materials should be addressed to Lars Steinmetz.

## Supplemental Figures

**Fig. S1:**
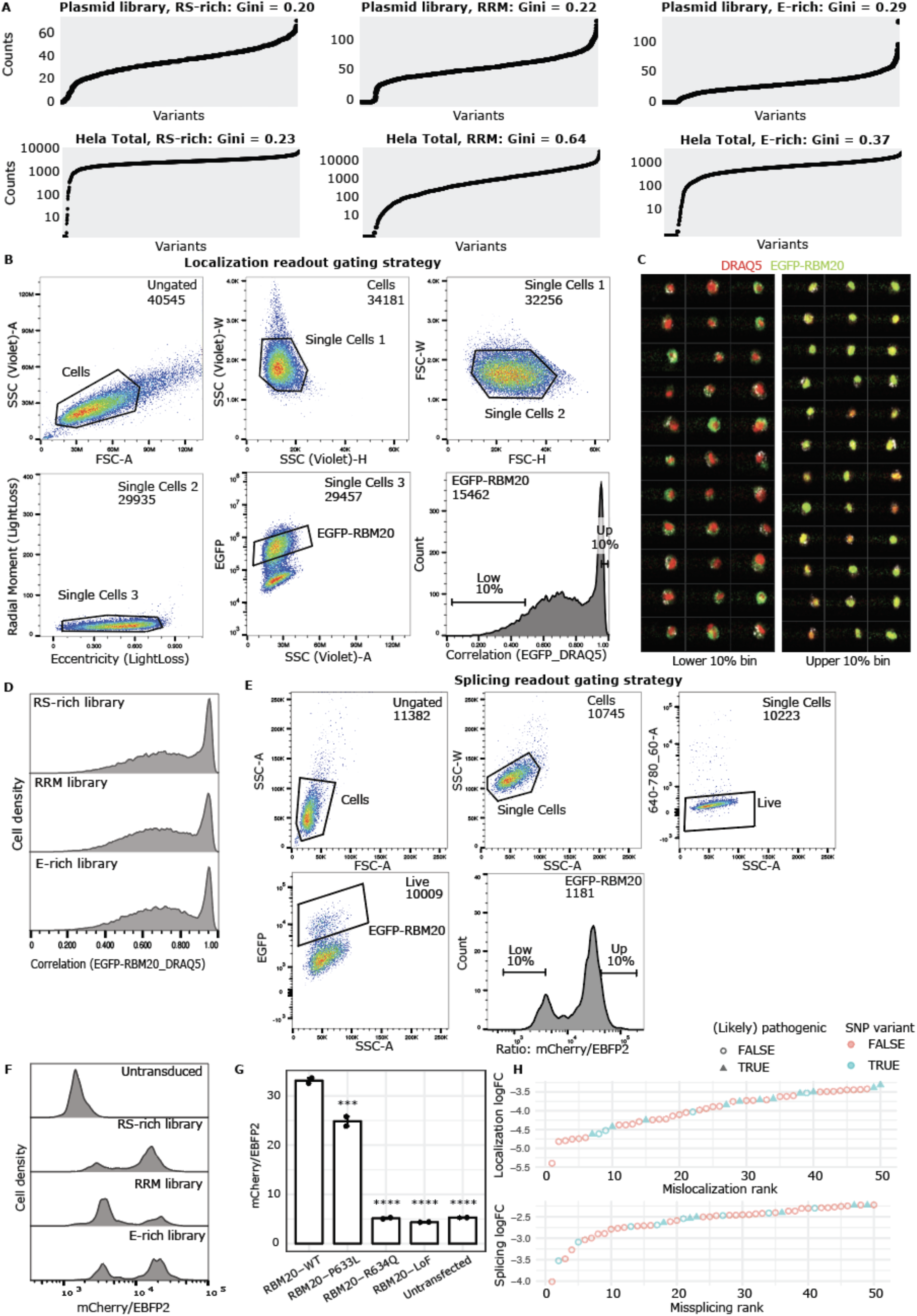
Multimodal phenotyping framework of localization and splicing activity. **(A)** Representation of single amino acid substitution variants in plasmid libraries and HeLa Kyoto cell pools for each screened domain. **(B)** Gating strategy for localization screen using Image-enabled Cell Sorting (ICS). **(C)** Example ICS images of cytoplasmic (lower 10% bin of EGFP:DRAQ5 correlation; left) and nuclear (upper 10% bin; right) sorted pools, at a sorting speed of 2,000 cells/sec. **(D)** ICS EGFP:DRAQ5 correlation plots for RS-rich, RRM, and E-rich mutagenesis HeLa Kyoto cell pools. **(E)** Gating strategy for splicing screening using FACS. **(F)** mCherry-to-EBFP2 distribution of HeLa Kyoto *Ttn* reporter cells transduced with each mutagenesis library. **(G)** Arrayed validation of the splicing readout strategy by transfection of HeLa Kyoto *Ttn* reporter cells with RBM20 variant constructs. Loss-of-function (LoF) variant refers to Ser635FrameShift^16^. *Ttn* N2A-and N2B-type isoform expression was assessed by flow cytometry. Mean ratios are reported; two biological replicates. Statistical significance was assessed with one-way two-sided ANOVA and Dunnett’s multiple comparisons vs. RBM20-WT: * - p < 0.05, ** - p < 0.01, *** - p < 0.001, **** - p < 0.0001. **(H)** Mislocalization and missplicing ranking of RS-rich domain variants (adj. p < 0.05; top 50 variants), turquoise color indicates variants that can occur via single nucleotide polymorphism and triangles known (likely) pathogenic variants (see **Table S5**).

**Fig. S2:**
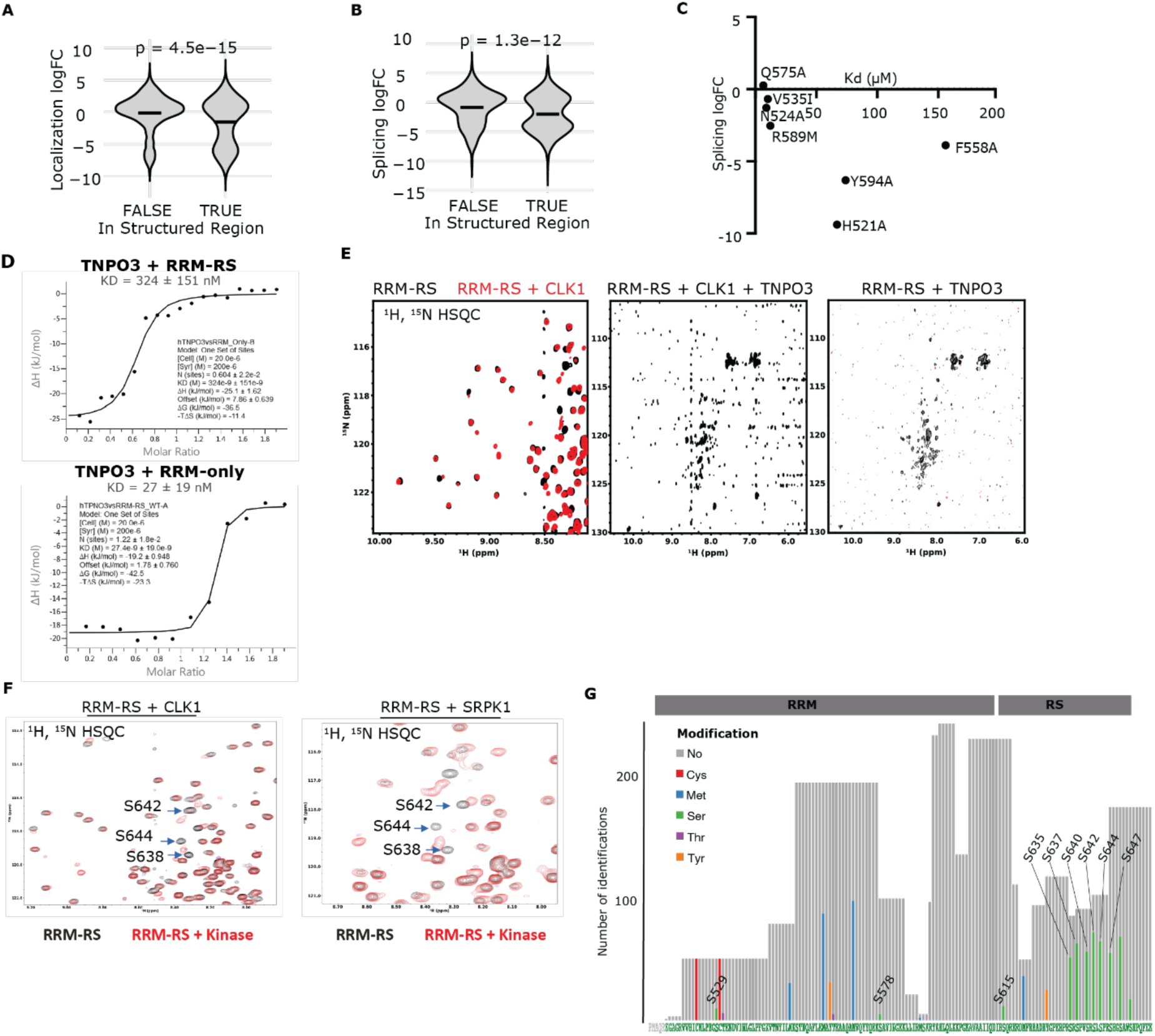
Impact of RRM-RS domain structure and phosphorylation on TNPO3 interaction. **(A-B)** Comparison of effect sizes of variants positioned within vs. outside annotated secondary structures. Statistical significance was assessed with a two-sided t-test. **(C)** Upadhyay & Mackereth^19^ isothermal titration calorimetry (ITC) measurements of RRM domain variants for mouse orthologs vs. functional splicing scores. Variants are labeled with human ortholog names. **(D)** Isothermal titration calorimetry (ITC) measurements of TNPO3 with RRM-RS wild type (left) or RRM alone (right), additional replicates. **(E)** NMR analysis: Left: Overlay of ^15^N,^1^H-HSQCs of unmodified RRM-RS (black) and phosphorylated after addition of CLK1 (red). Middle: ^15^N,^1^H-HSQC of the same sample after addition of TNPO3 (1:1) and CLK. Right: ^15^N,^1^H-HSQC of the same sample after addition of CLK1 first to RRM-RS and then TNPO3 (1:1). The disappearance of all resonances of the RRM and RS-rich domains confirms binding of TNPO3 to phosphorylated and non-phosphorylated RBM20-RRM-RS. **(F)** NMR validation of *in vitro* phosphorylation of the RRM-RS-rich domain. **(G)** Mass spectrometry results of *in vivo* CLK1 phosphorylated RRM-RS.

**Fig. S3:**
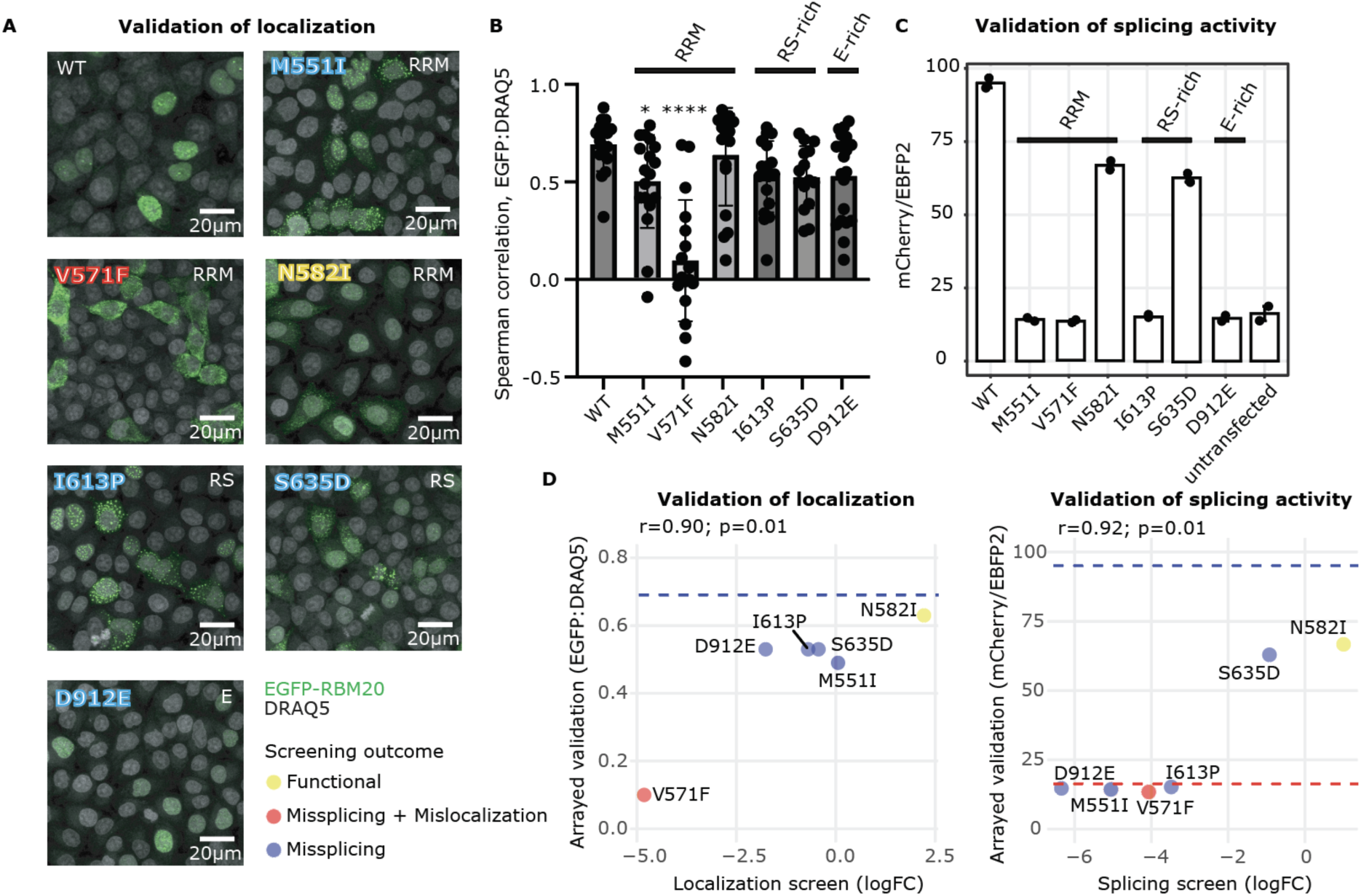
Functional validation of single variants. **(A)** Representative fluorescence microscopy images of HeLa Kyoto cells transfected with RBM20 variant constructs for arrayed validation of single variants. Overlays of EGFP (RBM20) and DRAQ5 (nucleus) channels are shown. Yellow (functional), red (missplicing + mislocalization), and blue (missplicing only) labels indicate the functional outcome based on our screen (threshold of adj. p < 0.05). **(B)** Quantification of EGFP and DRAQ5 signals using Spearman correlation. At least 15 regions of interest per variant were quantified via Coloc 2 analysis using ImageJ2 v2.14.0/1.54q. Statistical significance was assessed with one-way two-sided ANOVA and Dunnett’s multiple comparisons vs. WT: * - p < 0.05, ** - p < 0.01, *** - p < 0.001, **** - p < 0.0001. **(C)** Arrayed splicing validation of single variants by transfection of HeLa Kyoto *Ttn* reporter cells with RBM20 variant constructs. *Ttn* N2A- and N2B-type isoform expression was assessed by flow cytometry. Mean ratios are reported; two biological replicates. **(D)** Pearson correlation analyses of validation and screen effect sizes. The blue dashed line represents the RBM20-WT reference level. The red dashed line indicates the splicing activity level without RBM20 (untransfected control). Yellow (functional), red (missplicing + mislocalization), and blue (missplicing only) dots indicate the functional outcome based on our screen (threshold of adj. p < 0.05).

**Fig. S4:**
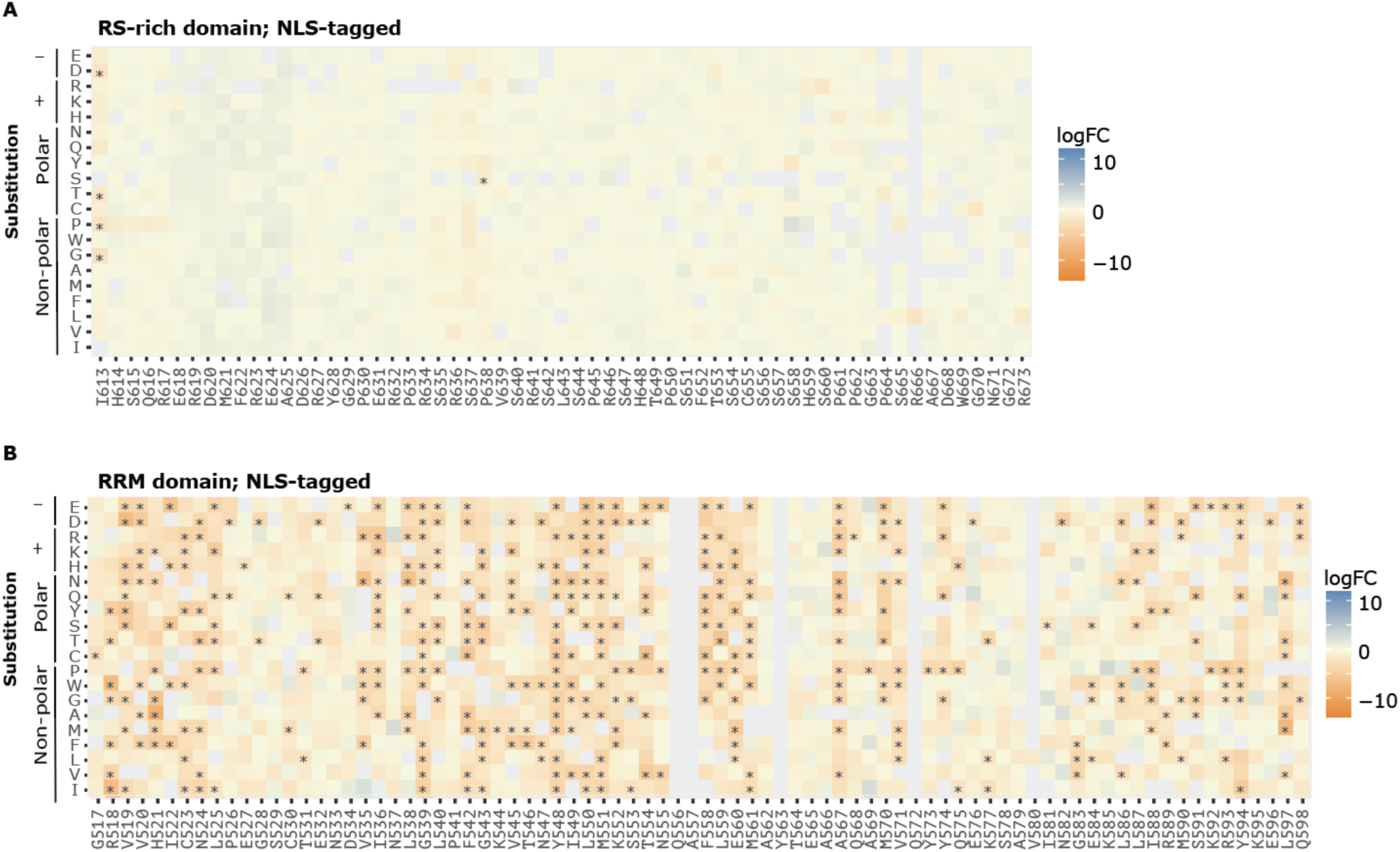
Systematic evaluation of nuclear re-localization. **(A-B)** Deep mutational scanning results for the NLS-tagged RS-rich and RRM domain mutagenesis library depicted as heat maps. Gray tiles indicate variants with missing data or WT residues. Heatmaps show results from at least two replicates, stars indicate significant single variant hits (adj. p < 0.05). Effect sizes were normalized to 10 benign-predicted variants (see materials and methods).

**Fig. S5:**
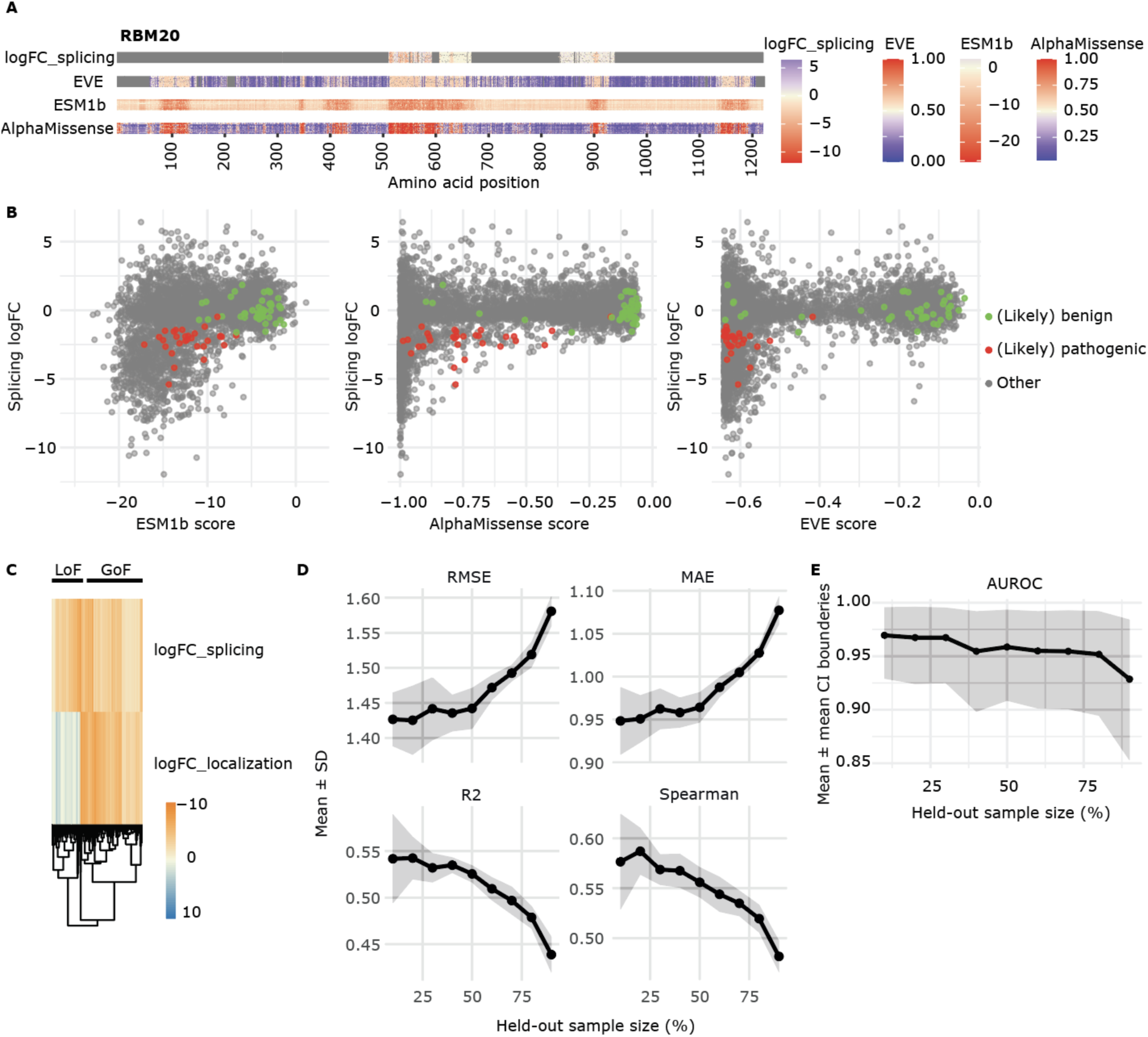
Comparison of functional scores to *in silico* predictions. **(A)** Our functional splicing scores compared to missense variant prediction maps of EVE, ESM1b, and AlphaMissense for the full length of human RBM20. Red indicates pathogenicity; blue indicates benignity. **(B)** Comparison of functional splicing scores and VEP scores. Annotated RBM20 variants are labeled (**Table S5**). The directionality of VEP scores was adjusted where applicable (multiplied by -1) so that smaller values indicate pathogenicity. **(C)** Identified gain-of-function (GoF) variants (logFC < -0.5 & adj. p < 0.05 for splicing and localization) and loss-of-function (LoF) variants (logFC < -0.5 & adj. p < 0.05 for splicing & logFC >= 0 for localization) visualized in a heatmap applying Euclidean distance for clustering and “average” as the clustering method. **(D-E)** Random forest models were trained to predict splicing logFCs using randomly sampled variant subsets from 10% to 90%. For each level, the variant training set was randomly sampled 10 times with different seeds. Reported values are the mean and standard deviation across repetitions. Model performance was evaluated using root mean squared error (RMSE), mean absolute error (MAE), coefficient of determination (R²), and Spearman correlations between measured and predicted values. Bootstrapping with 1000 iterations was performed to estimate AUROC with confidence intervals (CI) on held-out annotated variants (E).

**Fig. S6:**
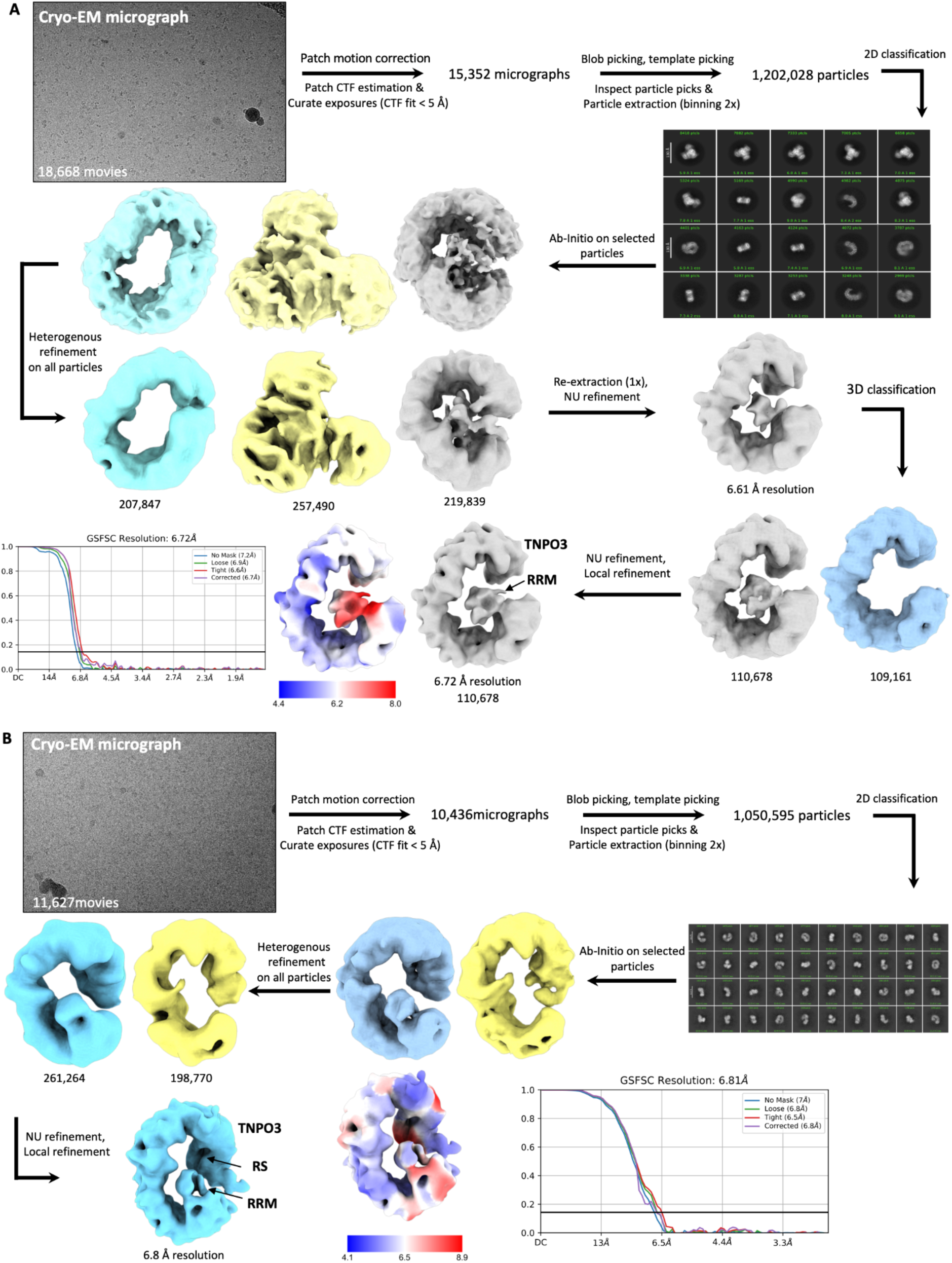
Cryo-EM workflow, data collection and refinement. **(A)** Cryo-EM workflow for TNPO3/RRM-RS in the unphosphorylated state **(B)** Cryo-EM workflow for TNPO3/RRM-RS in the phosphorylated state

## Materials and Methods

### Mutagenesis library design

Mutagenesis libraries for the three RBM20 domains (RS-rich, RRM, E-rich) were separately ordered from TwistBioScience (**Table S7**). Each library was designed to substitute every residue with each of the 19 possible amino acid alternatives in equal proportions. Stop codons and deletions were excluded. Codon usage was set to Homo sapiens, and restriction sites BsrGI, XbaI, and Esp3I were excluded.

RS-rich library: 61 positions (aa 613–673) were targeted; 60 passed quality control (QC). Position Arg666 and six single variants (S642D, P661C/G/S, S665R, D668C) were absent (**Table S7**). Final size: 1,134 variants (expected 1,159).

RRM library: 82 positions (aa 517–598) were targeted; 6 failed QC (Q556, A557, Y563, Q572, V580, plus variant E565W) (**Table S7**). Final size: 1,459 variants (expected 1,558).

E-rich library: 107 positions (aa 839–945) were targeted; 8 failed QC (D839, D840, R841, Q869, E870, E874, P878, Q937). Additionally, G897 lacked all variants except W, V, and E, and two variants (S862C, E944H) were missing (**Table S7**). Final size: 1,863 variants (expected 2,033).

### Mutagenesis cloning

All libraries were shipped in 96-well plates with 50 ng DNA per well with all variants of one position per well. To each well, 50 μl water was added and 2 μl of each well were mixed. The following primers were used to amplify the respective RBM20 domain, adding adjacent Esp3I restriction sites: RS-rich (KF069 + KF070), RRM (KF082 + KF083), E-rich (KF085 + KF086). All primers used in this study are listed in **Table S8**. The following PCR program was used: 95°C 2 min / 12 cycles (98°C, 20 sec/ 60°C, 20 sec/ 72°C, 30 min)/ 72°C 5 min. Each library was purified with PCR clean-up beads according to the provider’s manual (CleanPCR, CleanNA).

A lentiviral vector encoding human *RBM20* (*TetO-EGFP-GGSG-FLAG-RBM20* ^7^) was used to create three plasmids, by exchanging the RS-rich, RRM, or E-rich domain by a gBlock (ordered from IDT; sequences listed in **Table S8**) that integrates an Esp3I landing pad. This was done by double digest of the lentivirus targeting plasmid with either PasI and SbfI or PasI and BsaBI, and the respective gBlock was cloned in via Gibson assembly. Plasmids were confirmed by Sanger sequencing before use.

A 3x molar excess of purified library PCR products was used with the respective lentiviral backbone. The Golden Gate assembly reaction consisted of T4 DNA ligase buffer with 10 mM ATP (NEB), Esp3I (NEB) and T4 DNA ligase (NEB). The reaction mix was incubated in a thermocycler with the following settings: 25 cycles of: 37°C, 5 min + 16°C, 5 min and at the end 1h incubation at 37°C. A plasmid safe (Lucigen) reaction was performed by incubation at 37°C for 2h in the presence of additionally added Esp3I to ensure that all religated or unligated plasmids are degraded. The reaction was heat-inactivated at 70°C for 20 min and used for heat shock transformation of NEB 10-beta cells. The transformed bacteria were plated on eight 245 x 245 mm LB + ampicillin plates. The cells were harvested by adding LB media onto the plates and scraping all cells off. The cell suspensions were combined in 50 ml falcon tubes and centrifuged at 4,000 rpm for 10 min. The pellets were frozen in liquid nitrogen and stored at - 20°C. Pellets were thawed on ice and used for plasmid purification according to the general provided manual (ZymoPURE II plasmid maxiprep kit).

### Cell culture maintenance

HeLa Kyoto cells (previously described in^35^) were maintained at 37°C and 5% CO2 in DMEM, 4.5 g/L high glucose without pyruvate (Gibco, 11965084) supplemented with 10% FBS Supreme (Pan Biotech, P30-3031), 1% Sodium Pyruvate (Gibco, 11360070), and 1% Penicillin-Streptomycin (Gibco, 15140122). For splitting, cells were washed once with PBS, and afterwards detached with 0.25% Trypsin-EDTA (Gibco, 25200056) for 5 minutes at 37°C. Detached cells were resuspended in fresh media, diluted to the desired concentration, and plated for cell maintenance. Cryopreservation of cells was done in the culture media supplemented with 10% DMSO.

HEK293FT cells (Thermo Fisher Scientific) were maintained at 37°C and 5% CO2 in DMEM, 4.5 g/L high glucose without pyruvate (Gibco, 11965084) supplemented with 10% FBS Supreme (Pan Biotech, P30-3031), 1% Sodium Pyruvate (Gibco, 11360070), 1% Penicillin-Streptomycin (Gibco, 15140122), 1x GlutaMax (Gibco, 35050061), 1x NEAA (Gibco, 11140050), and 0.5 mg/ml G418 (InvivoGen). For splitting, cells were detached with 0.05% Trypsin-EDTA (Gibco, 25300054) for 5 minutes at 37°C. Detached cells were resuspended in fresh media, diluted to the desired concentration, and plated for cell maintenance. Cryopreservation of cells was done in the culture media supplemented with 10% DMSO.

All cell lines were regularly checked to be not infected with mycoplasma (via mycoplasma testing service of Eurofins).

### Lentivirus production

For lentivirus production, one 6-well plate of ∼90% confluent HEK293FT cells was used. First, the plasmid library was mixed with two lentivirus packaging vectors (pMD2.G, psPAX2), at equal ratios (total of 2,5 μg of DNA / transfected well of a 6-well plate). The solution was combined with Lipofectamine 3000 transfection reagent (Thermo Fisher Scientific), incubated at room temperature for 15 min, and added to HEK293FT cells. 6 h post-transfection, cells from each well were detached and cells from two wells were transferred into one 15 cm dish and incubated at 37°C and 5% CO2 for 3 days. After incubation, the cell culture supernatant was collected into a Falcon tube and centrifuged at 500 x g and 4°C for 10 min. The clarified supernatant was transferred to a new tube, mixed with Lenti-X Concentrator (Takara) at 3:1 ratio, and incubated at 4°C overnight. Next, the samples were centrifuged at 1500 x g and at 4°C for 45 min. The supernatant was removed, and the pellet was slowly overlaid with 500 μl PBS (to remove any remains of media) which was again aspirated. The pellet was then resuspended in 900 μl of cold PBS, and stored in 100 - 200 μl aliquots at -70°C.

### Lentivirus transduction efficiency test

The lentivirus transduction efficiency test was done with each new batch of lentivirus to determine the appropriate volume of lentivirus per transduction. The day before transduction, 0.18 million live HeLa Kyoto cells were seeded per well of a 6-well plate to reach 40-50% confluency after 24h. On the day of transduction, varied amounts of lentivirus (0, 1 µl, 5 µl, 10 µl, 25 µl or 50 µl per well) were added to a total volume of 1.3 ml culture media supplemented with Polybrene transfection reagent (Milipore, final concentration 10 µg/ml). 24h after the transduction, cells were detached and re-plated in 2 ug/ml Puromycin (Gibco, A1113803). Cell survival was monitored for 3 days, and the lentivirus volume that resulted in 5-10% survival was chosen for screening (proportionally upscaled to the surface of a 15 cm dish).

### Large-scale transduction for screening

One day prior to transduction, 3 million live HeLa Kyoto cells were seeded per 15 cm dish to reach 40-50% confluency after 24h. The day of viral transduction, 20 ml of fresh culture media was supplemented with Polybrene (Milipore, final concentration 10 µg/ml) and the previously tested volume of lentivirus to obtain 5-10% surviving cells. The number of plates per library was adjusted to aim for a 1000x coverage of library elements. 24h after transduction, HeLa Kyoto cells were detached and re-plated in fresh culture media supplemented with 2 μg/ml Puromycin (Gibco, A1113803). The cells were treated with Puromycin for 2-4 days, until the untransduced control condition showed no surviving cells. Afterwards, transduced HeLa Kyoto cells were cultured and expanded in normal culture media. Cells were harvested for assays no earlier than 6 days post-transduction.

### Harvest and fixation

HeLa Kyoto cells were harvested and fixed using 3% Glyoxal solution (freshly prepared on day of use: 3% glyoxal, 0.75% Acetic Acid-Glacial, 140 mM NaCl, pH = 5). Transduced and puromycin-selected HeLa Kyoto cell pools were detached using 0.25% Trypsin (Gibco), centrifuged at 500 x g and 4°C for 5 min, and resuspended in PBS to obtain approx. 10 million cells/ml. Next, 1 ml of cells were centrifuged at 500 x g and 4°C for 5 min, resuspended in 2,5 ml of 3% Glyoxal solution, and incubated at room temperature for 15 min. After incubation, 10 ml of ice-cold 2% BSA (Sigma) in PBS was added and the suspension was centrifuged at 500 x g and 4°C for 3 min. The washing step was repeated, and the cells were resuspended in ice-cold PBS (∼10 million cells/ml). Cell suspensions were filtered through 100 µm pluriStrainer mesh (Corning) and stored at 4°C until cell sorting.

### Localization screen using ICS

ICS was performed using the BD CellView^TM^ Imaging Technology^14^ on an S8 Discovery with a 100 μm sorting nozzle. The setup of an RBM20 localization screen was previously described for the purpose of a CRISPR-knockout screen^7^. Before the sort, fixed cells were stained with 1:60 v/v DRAQ5 (BioStatus, 100 mM final). Single cells that were DRAQ5 and EGFP positive were gated, and a histogram of the correlation of EGFP-RBM20 signal and nuclear DRAQ5 signal was plotted. The upper and lower 10% of the histogram were selected for bulk sorts of 100,000 cells. Also, 100,000 cells were sorted using the parental gate (input). All sorts were performed in purity mode. ICS data were analyzed using the FlowJo_v10.7.1_CL software. Sorted cells in 1.5 ml Eppendorf tubes were centrifuged at 10,000 x g and 4°C for 2 min, and the supernatant was discarded. The cell pellets were stored at -70°C until genomic DNA extraction.

### Splicing screen using FACS

We used a fluorescent *Ttn* mini gene compatible with our EGFP-RBM20 variant library by adapting the previously generated *TtnE50-E51E218-E219-EGFP/mCherry* construct^15^ containing the indicated murine *Ttn* exons (kindly provided by the Kuroyanagi lab): EGFP was changed to EBFP2 by Gibson Assembly (primers LM161 and LM162 used with original plasmid; primers KF138 and KF139 used with Addgene plasmid #194326 for EBFP2 amplification; XhoI and Esp3I digest of amplified products and the original plasmid), and subsequent site-directed mutagenesis of an obtained clone using primers LM163 and LM164 with the KOD Hot Start DNA Polymerase (Sigma) and the Q5 Site-Directed Mutagenesis Kit (NEB). The resulting *TtnE50-E51E218-E219-EBFP2/mCherry* mini gene was further cloned into a vector with PiggyBac inverted terminal repeat sequences. A reporter cell line was generated by co-transfection of HeLa Kyoto cells with the PiggyBac *Ttn* mini gene construct and a hyperactive PiggyBac transposase encoding plasmid^35^, kindly provided by the Craig lab. 10 days after transfection, single positive clones were isolated via FACS. Clones were expanded and, 12 days after sorting, again screened for stable expression of the mini gene via flow cytometry.

Transduced and selected HeLa Kyoto *Ttn* reporter cells were harvested, stained with LIVE/DEAD Fixable Near-IR Dead Cell dye (Invitrogen), and preserved via glyoxal fixation as described above. FACS was performed on the BD FACSAria Fusion Flow Cytometer with a 100 μm sorting nozzle. The gating strategy filtered cell debris, doublets, dead cells, and EGFP-RBM20-negative cells. Upper and lower 0-10% and 10-20% of the mCherry:EBFP2 distribution were sorted, with a coverage of 100x cells per library element. Also, input samples were sorted using the parental gate. All sorts were performed in purity mode. Flow cytometry data were analyzed using the FlowJo_v10.7.1_CL software. Sorted cells in 1.5 ml Eppendorf tubes were centrifuged at 500 x g and 5°C for 8 min, and the supernatant was discarded. The cell pellets were stored at -70°C until genomic DNA (gDNA) extraction.

### Genomic DNA extraction + amplicon sequencing

The cell pellets were thawed on ice, 100 μl ice cold PBS was added, and each pellet was resuspended. The genomic DNA was extracted according to the provided manual using the Monarch genomic DNA purification kit (NEB). Elution was done twice using 35 µl elution buffer. The eluted genomic DNA was measured by Nanodrop and the total DNA was used for Amplicon Sequencing. Three primer pairs were used for amplification: KF038 + KF039 for the RS-rich domain, KF116 + KF117 for the RRM domain, and KF118 + KF119 for the E-rich domain. Staggered primers of these were used that added 1 to 3 nucleotides before the amplicon. PCR1 was performed with the 2X Q5 Hot Start High-Fidelity Master Mix (NEB) and run with the following thermocycler program: 5 min 95°C / 20x (30 sec 95°C, 20 sec 65°C, 30 sec 72°C)/ 1 min 72°C. For RRM domain samples, Q5 High GC Enhancer was added to the PCR1 reaction. Without further purification, 2 μl of each PCR1 reaction was transferred to PCR2 that added the Illumina overhangs (Nextflex primers), also using the 2X Q5 Hot Start High-Fidelity Master Mix (NEB) and the following thermocycler program: 1 min 98°C / 12x (10 sec 98°C, 20 sec 65°C, 20 sec 72°C)/ 2 min 72°C. PCR2 was purified with 1.2x (RS-rich and E-rich) or 0.7x (RRM) AMPure XP beads (Beckman) according to the provided manual, and DNA concentrations were measured by Qubit using the dsDNA High Sensitivity kit (Invitrogen). Samples were run on a Bioanalyzer High Sensitivity DNA chip (Agilent) to confirm correct amplicon size (expected size PCR1/PCR2: RS-rich = 334/402 nt, RRM = 343/411 nt, E-rich = 428/496 nt). A multiplex was generated and sequenced on a MiSeq or NextSeq2000 Illumina sequencer, aiming for a minimum of 200 reads per library element per sample.

### Screen data analysis and visualization

Read 1 and read 2 were merged per sample using FLASH^36^. Merged reads were analyzed using the mutscan R package^37^ (v0.3.4). Variant calling was performed by alignment to the RBM20 WT sequence. Reads with multiple mutated codons were removed. Variant counts were grouped by amino acid exchange. Variants with a total read count < 100 or an individually chosen threshold above (relying on sequencing depth), and variants appearing only in a single sample were excluded from analysis. LogFCs and associated adjusted p-values were calculated for each variant using the limma setting. Effect sizes were normalized, per domain, to the mean of the AlphaMissense^26^ top-10 most benign-scored variants among those detected in all three readouts (splicing, localization; bottom quartile excluded). These include “Met621Leu”, “Ala625Thr”, “Met621Val”, “Pro630Thr”, “Ala625Val”, “Ala667Thr”, “Gly672Ser”, “Pro630Leu”, “Ala625Ser”, and “Gly663Ser” for the RS-rich domain; “Ser578Ala”, “Ser578Pro”, “Asn582Ser”, “Ser578Thr”, “Gln568Glu”, “Lys577Arg”, “Ser578Cys”, “Gln575Glu”, “Gln568Lys”, and “Asn537Ser” for the RRM domain; and “Glu892Gly”, “Thr845Arg”, “Phe875Ser”, “Phe875Ile”, “Met846Val”, “Thr845Lys”, “Gln864Pro”, “Phe875Val”, “Ala873Thr”, and “Phe875Thr” for the E-rich domain. Hits were defined as adjusted p-value < 0.05. Effect sizes of variants were visualized in heat maps using ggplot2 (v3.5.2).

For collapsed plots of fold stability predictions^17^, amino acids were grouped by chemical and biophysical properties and the average effect was calculated per group. These groups are non-mutually exclusive. Non-polar residues include A, V, L, I, M, F, W, P, G; Polar residues include S, T, N, Q, Y, C; Positively-charged residues include K, R, H; Negatively-charged residues include D, E. Ring residues include F, Y, W, H, P; Non-bulky residues include A, M, G, S, T, N, Q, C, K, R, E, D; Branched residues include V, I, L.

### RBM20 variant annotations

RBM20 variant annotations used in this study originate from the ClinVar database^31^ (accessed 17/07/2025) and internal curation. Variants were classified by a board-certified medical geneticist based on American College of Medical Genetics and Genomics criteria, with modifications for DCM^38^. To maximize the number of variants available for screen validation, we defined a variant as (likely) pathogenic or (likely) benign in cases where at least one source classified it as such. Thereby, the annotation set included a total of 49 (likely) benign, 30 (likely) pathogenic, and 154 VUS/Conflicting missense variants across the screened domains (**Table S5**).

### Random forest modeling

Random forest models were trained to predict splicing effect sizes from a set of 46 features (**Table S6**) and localization DMS scores (effect sizes and adjusted p-values). For each run, a random test set was selected by sampling a fixed proportion of variants (10 to 90%, in 10% steps). A random forest was fitted on the remaining training data using the R package randomForest (v4.7-1.2). The model was trained with the randomForest() function, specifying importance = TRUE and ntree = 100. Predictions were generated for the masked test set. Predictive performance was evaluated by RMSE, MAE, R², and Pearson/Spearman correlations between predicted and observed splicing effect sizes. Predicted scores were merged with RBM20 variant annotations, mapping labels to a binary class ((Likely) pathogenic = 1; (Likely) benign = 0). To assess variability, the random hold-out procedure was repeated 10 times per test set size.

The AUROC was computed with the pROC package (v1.18.5) using the roc() function. Confidence intervals were obtained with the ci.auc() function based on 1,000 bootstrap replicates (method = “bootstrap”, conf.level = 0.95), where each replicate sampled 100% of the original data with replacement. When using the AUROC as an evaluation metric, ClinVar annotations were excluded from training features to prevent circularity, and annotated variants (**Table S5**) were held out for testing.

### Single variant validations

The day before transfection, 10,000 live HeLa Kyoto *Ttn* reporter cells were seeded per well of a 96-well plate. Transfection was done in arrayed fashion using *EGFP-RBM20* cDNA variant constructs^7^, 0.3 µl Lipofectamine 3000 (Thermo Fisher Scientific), and 100 ng plasmid DNA per well.

48 h to 72 h after transfection, cells were analyzed on a BD FACSymphony A3 Cell Analyzer. DRAQ7 (Thermo Fisher Scientific) staining was performed prior to analysis to exclude dead cells. The gating strategy further filtered cell debris, doublets, and EGFP-negative cells. The RBM20 splicing activity was assessed by the mCherry-to-EBFP2 signal ratio of the gated population. Flow cytometry data were analyzed using the FlowJo_v10.7.1_CL software.

72 h after transfection, cells cultured in glass-bottom plates were washed with PBS and fixed with 4 % PFA in PBS. DRAQ5 (Biostatus, 1:500 dilution in PBS) staining was performed prior to acquisition of fluorescence microscopy images. Confocal microscopy was performed as previously described^7^. The maximum-intensity projections of the Z-stacks with modified intensities were used for quantification. The EGFP-RBM20 and DRAQ5 signal correlation was calculated with the Coloc 2 plugin of ImageJ2 v2.14.0/1.54q. At least 15 regions of interest with EGFP-positive cells per variant were quantified, and Pearson and Spearman correlation coefficients were reported.

### RRM-RS protein purification

Different RRM-RS constructs were initially cloned including different length of the RS-rich domain (full-length RRM-RS: residues 511-673, truncated #1 RRM-RS: 511-668, and truncated #2 RRM-RS: 511-648). The best solubility was observed for the second truncated construct and was used for all downstream protein purifications as pETM-14-RBM20 (511-648, pETM-14 backbone from Novagen) wild type or single residue variants R634Q or P633L. In addition, one RRM-only construct (residues 511-594) was generated and purified as described below. The plasmid was transformed into BL21-CodonPlus-RIL (Agilent) competent cells and overnight cultures were grown in LB media supplemented with 1% glucose and 2 mM MgSO_4_, as well as 30 µg/ml Kanamycin and 34 µg/ml Chloramphenicol. After 16 hours of incubation, 1L of TB media with the same supplements as LB and 1.5% lactose was inoculated and incubated at 37°C and 200 rpm in a non-baffled flask. At an OD of 0.8, the flasks were transferred to 18°C and induced with 0.05 M IPTG for 18 hours. Dense cultures were subsequently centrifuged in a JLA8.100 rotor at 4,000 rpm and room temperature for 25 min, and the cell pellet was frozen in liquid nitrogen and stored at -20°C. Pellets were thawed on ice and resuspended in 20 ml high salt lysis buffer (50 mM Tris pH 7.5, 500 mM NaCl, 20 mM Imidazole) on ice. One protease inhibitor tablet (Roche, Complete EDTA free protease inhibitor) was added per 20 ml lysate. Cells were lysed with a Microfluidizer LM10 (Microfluidics) with 4 repeats at a pressure of 12,000 psi. Lysates were centrifuged at 50,000 x g and 4°C for 30 min in a JA 25.50 rotor. Supernatant (soluble fraction) was added to a column with Ni-NTA slurry (50% solution, MACHEREY-NAGEL, Protino Ni-NTA) equilibrated with high salt lysis buffer at 4°C (two times). The column was washed with a high salt lysis buffer and then eluted with an elution buffer (with 25 mM Tris pH 7.5, 250 mM NaCl, 250 mM imidazole). To 7 ml elution sample, 385 µl of 3C-protease (1 mg/ml) was added and incubated overnight at 4°C. The 3C-cleaved sample was directly loaded onto a StrepTactin slurry containing column (50% solution, IBA Strep-Tactin Superflow high capacity) and the flowthrough was loaded a second time to increase the fraction of bound protein. A wash step with a low salt buffer was performed (25 mM Tris pH 7.5, 150 mM NaCl). The sample was eluted in 20 mM Tris-HCl pH 8, 150 mM NaCl, 10 mM Desthiobiotin and fractionated in 1 mL fractions. All protein-containing fractions were loaded on an equilibrated Superdex S200 column (20 mM NaH_2_PO_4_/Na_2_HPO_4_ at pH 6.5, 100 mM KCl, 1 mM DTT). Protein-containing fractions were combined and added to a concentrator tube with a 3 kDa cutoff (Amicon Ultra, Merck) and centrifuged at 4,000 rpm at 4°C. Protein samples were concentrated to 5 mg/ml and then frozen in 50 μl fractions in liquid nitrogen and stored at -80°C. The same procedure was applied for ^15^N or ^15^N, ^13^C labeled protein samples with the change of growth media, which contained 1x M9 salt solution without NH_4_Cl, (for ^15^N labelling) 0.4% Glucose, 1 mM MgSO_4_, 1 µg biotin, 1 µg thiamin, 1 µg riboflavin, 0.5 g/l ^15^NH_4_Cl, 2 g/l ^13^C-Glucose (for ^15^N, ^13^C labelling), and the respective antibiotics.

To obtain the phosphorylated RRM-RS sample, we performed a co-transfection of the same pETM-14-RBM20 purification plasmid and pLIC-SGC-CLK1 (Addgene, #39149). E.coli cells were cultured as described above with the respective antibiotics. The purification was conducted as described for the non-phosphorylated version with the addition of a further purification by anion exchange chromatography on a 5-ml HiTrap Q sepharose column (GEHealthcare) and gel filtration through a HiLoad 16/60 Superdex 200 column in 300 mM NaCl, 25 mM Tris-HCl, pH 7.4. Peak fractions of phosphorylated RRM-RS proteins were supplemented with 2 mM DTT and 10% glycerol and stored at -80°C.

### TNPO3 protein purification

The full-length human TNPO3 cDNA sequence was cloned into the pET24-his-Sumo-hTNPO3 plasmid. TNPO3 was purified as above, with the following exceptions. The lysis buffer was exchanged against: 25 mM Tris-HCl, pH 8.0, 500 mM NaCl, 20 mM Imidazole, 10% glycerol, Protease Inhibitors (cOmplete Roche). After the Ni-NTA purification, the protein sample was treated with the SUMO specific peptidase SENP2 and dialyzed overnight in lysis buffer and 20 mM imidazole prior to further purification. Samples were loaded onto a reverse Ni-Q combo column to remove the cleaved off His-SUMO tag. Protein containing samples were loaded for Size Exclusion Chromatography (SEC) on Superdex 200 16/600 at 1.0 ml/min in 25 mM Tris pH 7.4, 200 mM NaCl, and 10% glycerol. The column was washed and eluted in lysis buffer with 250 mM imidazole. Protein-containing fractions were pooled and added to a concentrator tube with a 30 kDa cutoff (Amicon Ultra, Merck) and centrifuged at 4,000 rpm at 4°C. Protein samples were concentrated to 10 mg/ml and then frozen in 50 μl fractions in liquid nitrogen and stored at -80°C.

### ITC measurements

One day before the measurement, all required protein samples (TNPO3 and RRM-RS constructs) were dialyzed in 25 mM Tris pH 7.5 and 150 mM NaCl at 4°C in dialysis membranes with a 2 kDa cut off (Slide-A-Lyzer MINI, ThermoFischer). Dialyzed protein concentration was determined and samples were diluted in dialysis buffer (20 µM TNPO3 (350 ul) and 200 µM RRM-RS (80 ul)). All samples were centrifuged at 10,000 x g for 5 min and then equilibrated at room temperature for 10 min in a metal block before measuring the samples with the MicroCal PEAQ ITC (Malvern Panalytical).

### Complex formation of TNPO3/RRM-RS

Affinity-purified TNPO3 and RRM-RS proteins were mixed at a 1:2.25 molar ratio and incubated on ice for 30 minutes to allow complex formation. The resulting TNPO3/RRM-RS complex was injected into a Superdex 200 (SEC S200) analytical size-exclusion chromatography column. Peak fractions containing the TNPO3/RRM-RS complex were pooled and concentrated to 1 mg/ml for single-particle cryo-electron microscopy (cryo-EM) analysis.

### Cryo-EM sample preparation

Quantifoil R 2/1, Cu 200 mesh carbon grids (Quantifoil) were glow-discharged at 25 mA for 120 seconds using a Glow Discharge Unit (GlowCube). Then, 5 μl of the purified TNPO3/RRM-RS complex was applied to the glow-discharged grids and incubated for 10 seconds. Grids were blotted for 2 seconds with a blotting force of -7 at 4 °C and 100% humidity, followed by plunge freezing into a 63:37 propane:ethane mixture using a Vitrobot Mark IV (Thermo Fisher Scientific).

### Data collection and processing

Vitrified grids were initially screened on an Artica 200 kV transmission electron microscope (TEM) (Thermo Fisher Scientific) equipped with an autoloader to identify optimal grid areas. Grids with uniform ice thickness and even particle distribution were selected for high-resolution data collection on a Titan Krios G3i microscope (Thermo Fisher Scientific) operating at 300 kV. Images were acquired using a K3 direct electron detector (Gatan) in counting mode, with a 20 eV energy filter slit width. A total of 18,668 movies were collected at a nominal magnification of 105,000x, corresponding to a calibrated pixel size of 0.85 Å. Each movie stack consisted of 50 frames, with a total electron dose of 56.5 e⁻/Å² and a defocus range of -0.5 to -2.5 μm. Automated data acquisition was performed using EPU software (Thermo Fisher Scientific).

Data processing was carried out in cryoSPARC v4.2. Movies were motion-corrected using Patch Motion Correction, and contrast transfer function (CTF) estimation was performed using Patch CTF. Micrographs were curated using a resolution cut-off of 5 Å. Particles were initially picked using the Blob Picker tool and subjected to 2D classification. High-quality classes were used as templates for further particle picking, followed by multiple rounds of 2D classification to clean the particle stack.

A total of 1,202,028 particles were selected for ab initio reconstruction to generate six initial models for the unphosphorylated state (**Fig. S6A**). These were subjected to heterogeneous refinement into twelve classes. Four classes with well-defined structural features were selected for non-uniform refinement. Two volumes, corresponding to 12.2% and 13.16% of particles, were identified as TNPO3 alone, with resolutions of 5.8 Å and 6.5 Å, respectively. One volume (17.16% of particles) corresponded to a TNPO3 dimer at 4.8 Å resolution, and another (10.2% of particles) represented TNPO3 in complex with the RRM domain at 6.72 Å resolution. For the phosphorylated state 1,050,595 particles were selected for ab initio reconstruction in the same way as for the phosphorylated state (**Fig. S6B**). Particles that represented TNPO3 in complex with the phosphorylated RRM-RS domain resulted in a resolution of 6.8 Å.

### Model building and refinement

To build the TNPO3/RRM-RS complex model, AlphaFold3 (AF3) was used to predict the structure based on the individual sequences of TNPO3 and the RRM-RS domain. However, the resulting AF3 model did not fit the cryo-EM density map. In a subsequent attempt, the sequences of TNPO3 and RRM-RS were fused into a single polypeptide chain and submitted to AlphaFold3 (AF3), which generated a model that showed a good fit to the cryo-EM map, with a predicted TM-score (pTM) of 0.77.

To validate this model, the gapTrick program was employed by randomly orienting the RRM-RS domain in the context of TNPO3. The resulting model, which matched the AF3 configuration, yielded high confidences with a predicted local distance difference test (pLDDT) score of 91.10 and a pTM score of 0.82. This model was further refined using Servalcat, implemented in the CCP-EM Doppio suite.

### NMR spectroscopy

NMR backbone assignment experiments of truncated #2 RRM-RS (residues 511-648) were collected on a Bruker Avance III 800 MHz spectrometer equipped with a cryoprobe. The experiments were collected at 298 K with samples containing 10% D_2_O as a locking agent. The NMR sample buffer consisted of 20 mM sodium phosphate (pH 6.5), 100 mM KCl, and 1 mM DTT and the protein had a concentration of 300 µM. An isotopically ^15^N/^13^C-labeled was obtained as described above. Chemical shift assignments were obtained using standard ^1^H-^13^C-^15^N correlation experiments^39^ with apodization weighted sampling to reduce data collection times^40^. These have been deposited under the BMRB accession code 53366. The raw data were processed and analyzed using NMRpipe^41^ and CARA (http://cara.nmr.ch).

For assessing the phosphorylation state of truncated #2 RRM-RS, NMR experiments were conducted at 298 K sample temperature on a Bruker 700 MHz Avance IIIHD NMR spectrometer equipped with a cryoprobe. Phosphorylation was detected by recording ^1^H, ^15^N HSQC spectra of ^15^N labeled samples (50 µM protein in 25 mM Tris, pH 7, 150 mM NaCl, 2.5 mM DTT 10 mM MgCl_2_, 2 mM ATP) before and after addition of 2 µl CLK1 or SRPK1. Peaks corresponding to serine residues shifted upon phosphorylation by kinases CLK1 and SRPK1. To test binding of TNPO3 to phosphorylated RRM-RS, an additional ^1^H, ^15^N HSQC TNPO3 was recorded after addition of TNPO3 in a 1:1 ratio to the NMR sample of CLK-1 phosphorylated RRM-RS.

